# An astrocytic AMPAR-dependent parity signal preserves synaptic strength

**DOI:** 10.64898/2026.10.08.757679

**Authors:** Nimrod Elazar, J. Russell Ravenel, Jinhu Kim, Trisha Vaidyanathan, Brenda C. Shields, Olivia Wirfel-Farenchak, Kunihiro Uryu, Shaun S.X. Lim, James M. Roach, Weizhan Zhang, Xiangyu Cao, Micah E. van Praag, Haidun Yan, Guoqiang Yu, Michael R. Tadross, Cagla Eroglu

## Abstract

Astrocytic processes closely associate with excitatory synapses and express AMPARs, positioning them to sense the same glutamate signals as neurons. Whether a mismatch between neuronal and astrocytic glutamate sensing affects synaptic physiology is unknown. Using AstroHTP, an extension of Drug Acutely Restricted by Tethering (DART) for cell-type-specific pharmacology, we found that acute astrocytic AMPAR inhibition spared neuronal AMPARs and basal transmission yet converted ongoing glutamatergic activity into a remarkably potent form of long-term depression (LTD) in mouse striatal projection neurons (SPNs). Unlike canonical presynaptic LTD, this form of LTD manifests postsynaptically, requires non-ionotropic NMDA receptor signaling, and arises when glutamate transmission occurs without a matching astrocytic signal. Astrocytic glutamate sensing through AMPARs, therefore, acts as a parity signal that verifies transmission and stabilizes active synapses.

## Main Text

In the striatum, afferents from cortex and thalamus provide excitatory glutamatergic input to spiny projection neurons (SPNs), which sense glutamate rapidly through AMPA-type glutamate receptors (AMPARs) (*1–3*). Astrocytic processes closely associate with these synapses (*4–7*) and also express AMPARs, positioning them to sense the same rapid glutamate signals. Yet whether a mismatch between neuronal and astrocytic glutamate sensing affects synaptic physiology remains unknown.

Previous work shows that astrocytic AMPARs are functional and capable of detecting physiologically relevant glutamatergic activity (*8–10*). Functional Ca²⁺-permeable AMPARs have also been identified on astrocytes, where they mediate intracellular calcium signaling (*9*). However, causal studies have been difficult to implement. Inducible deletion of GluA1 and GluA4 from cerebellar Bergmann glia alters synaptic coverage, Purkinje cell currents, and motor coordination (*10*). Yet because receptor loss unfolds over weeks, glial remodeling and compensatory changes in gene expression complicate a direct causal link to astrocytic AMPARs themselves. Conversely, pharmacology is acute but cannot isolate the function of astrocytic AMPARs from their more abundant neuronal counterparts.

To overcome these limitations, we adapted Drug Acutely Restricted by Tethering (DART), a cell-specific pharmacology platform (*11, 12*), to acutely antagonize endogenous AMPARs in striatal astrocytes. Using this approach, we found that acute astrocytic AMPAR inhibition spared neuronal AMPAR function and basal transmission but converted ongoing glutamatergic activity into a noncanonical postsynaptically expressed form of long-term depression (LTD). Thus, astrocytic AMPARs sense synaptic glutamate and determine whether active synapses retain their strength or undergo depression. We propose that astrocytic glutamate sensing acts as a parity signal that verifies the fidelity of synaptic transmission.

### Establishing astrocytic DART for cell-type-specific pharmacology in intact circuits

DART enables cell-specific pharmacology through two components: a cell-surface HaloTag Protein (HTP) expressed on cells of interest, and a drug linked by polyethylene glycol (PEG) to the HaloTag Ligand (HTL) (Fig. 1A). Covalent binding between HTP and HTL locally accumulates the tethered drug to concentrations substantially higher than ambient levels. The second-generation DART.2 platform achieves thousandfold enrichment of tethered drugs within minutes, enabling potent modulation of endogenous receptors in defined neuron types *in vivo* (*11, 12*). However, DART has not yet been extended to astrocytes.

**Figure 1.**
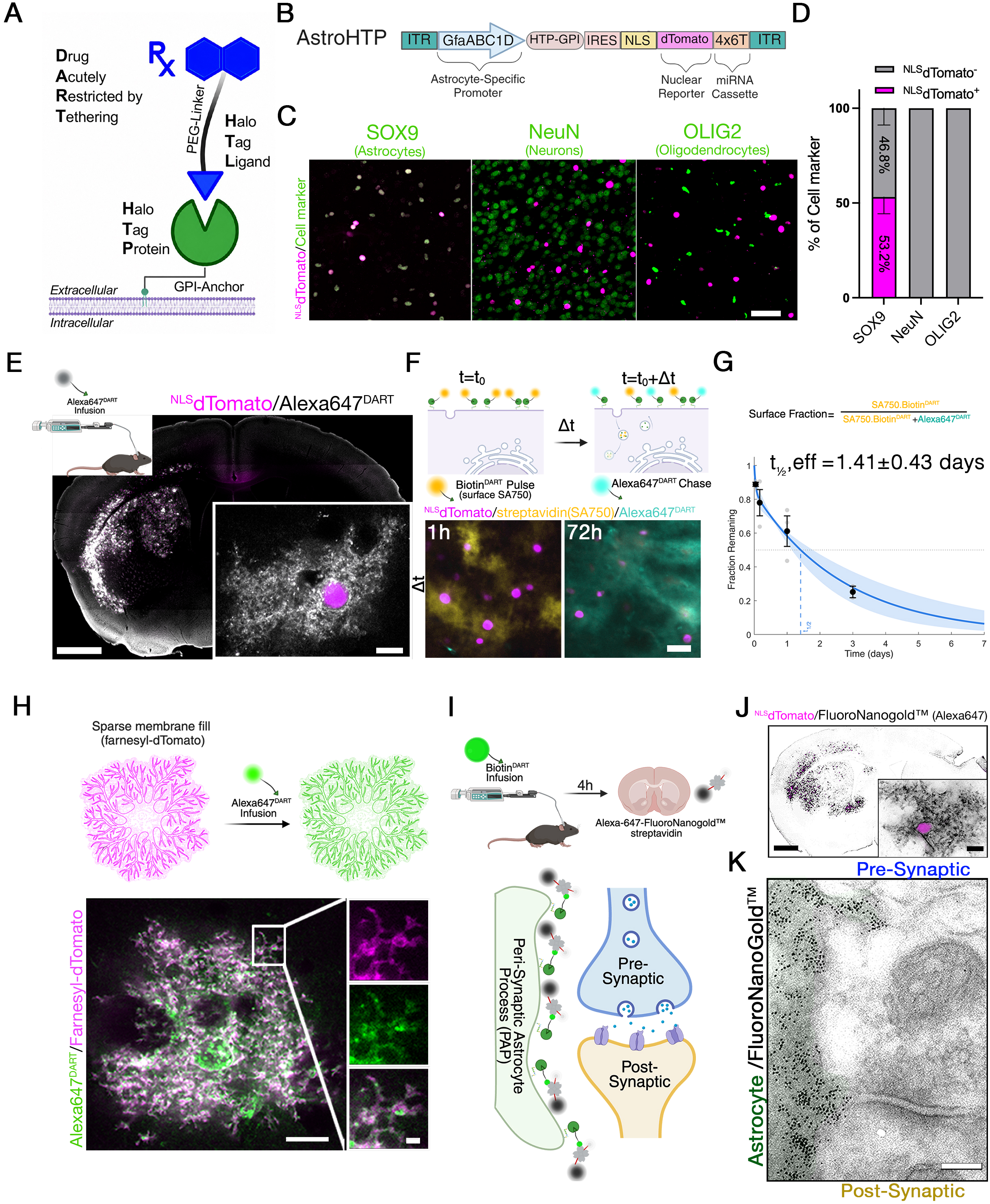
AstroHTP drives astrocyte-specific HaloTag expression and perisynaptic tethered-ligand capture in striatum. **(A)** Schematic of DART technology. Genetically specified cells express a GPI-anchored HaloTag Protein (HTP), which acutely tethers a drug (Rx) attached via a polyethylene glycol linker (PEG) to a HaloTag Ligand (HTL). **(B)** Schematic of the AstroHTP viral construct. HTP-GPI is expressed under control of the GfaABC1D promoter, followed by an IRES sequence and a dTomato reporter containing a nuclear localization signal (NLS). A 4×6T microRNA-targeting cassette is added to the 3’ UTR, and the construct is packaged in the PHP.AX capsid. **(C)** Three weeks post-AstroHTP injection, coronal mouse brain sections containing the dSTR were immunolabeled with antibodies against SOX9 (astrocytes), NeuN (neurons), and OLIG2 (oligodendrocytes). Scale bar, 100 µm. **(D)** Quantification of AstroHTP cell-type transduction efficiency, measured as the percentage of each cell type co-localized with dTomato-positive nuclei in the dSTR (N = 3 mice and 5 to 6 images per mouse). Data are mean ± SEM. **(E)** AstroHTP was injected into the dSTR of adult C57BL/6J mice. Three weeks later, 3 µM Alexa647 fluorophore conjugated to HTL (Alexa647^DART^) was infused into the striatum and histology was performed 24 h later. Representative coronal section, showing ligand capture (grey) and NLS-dTomato (magenta). Scale bars: 500 µm, 5 µm. **(F-G)** HTP turnover, **(F)** Experimental pulse-chase approach: a 0.9 µL pulse of 60 µM biotin-conjugated HTL (Biotin^DART^) was administered, followed 1–72 h later by a 0.9 µL chase of 60 µM Alexa647^DART^. Example histology at the 1 h and 72 h chase intervals. Scale bar, 20 µm. **(G)** Quantification of the fraction of original surface HTP signal remaining versus chase interval; each symbol represents an independent mouse, and error bars are mean ± SEM. Data were fit with a double exponential, yielding an effective half-turnover time for HTP of 1.41 ± 0.43 days (mean ± 95% CI, intervals derived from the 2.5th and 97.5th percentiles of the fitted parameter distribution). **(H)** Astrocytes capture DART ligands on their fine processes. A Cre-dependent HTP-GPI variant with a membrane-targeted reporter (dTomato-farnesylated) was co-injected with GfaABC1D-Cre into the dSTR of adult C57BL/6J mice. Three weeks later, 3 µM Alexa647^DART^ was infused into the dSTR of head-fixed mice, and histology was performed 24 h later. Representative coronal section showing DART ligand capture (green) and membrane-targeted dTomato (magenta). Scale bars: 10 µm; inset, 1 µm. **(I-K)** AstroHTP-transduced astrocytes capture DART ligands at perisynaptic processes. **(I)** AstroHTP was injected into the dSTR of adult C57BL/6J mice. Three weeks post-injection, 0.9 µL of 30 µM Biotin^DART^ was infused into the striatum, and histology was performed 4 h later. Coronal sections were processed for immuno-electron microscopy and labeled with Alexa647-conjugated FluoroNanogold™. Schematic of the tripartite synapse depicting perisynaptic Alexa647-conjugated FluoroNanogold™ capture **(J)** Representative coronal section showing immunohistochemistry for NLS-dTomato (magenta) and FluoroNanogold (black). Scale bars: 1000 µm, 10 µm. **(K)** A representative electron micrograph of the mouse striatum showing a nanogold-labeled perisynaptic astrocytic process (green). Scale bar, 100 nm.

We therefore developed AstroHTP, an AAV construct that achieves astrocyte–specific expression without the need for transgenic animals or recombinases. AstroHTP uses the astrocyte-specific GfaABC1D promoter (*13*) to express GPI-anchored HTP (HTP_GPI_) along with a bicistronic nuclear fluorescent reporter (IRES-^NLS^dTomato). To further enhance astrocyte specificity, we incorporated four repeats of a microRNA targeting cassette (4×6T), previously shown to suppress expression in neurons and endothelial cells (*14*) (Fig. 1B), and packaged the vector into the astrocyte-tropic PHP.AX capsid (*15*).

To assess expression penetrance and astrocyte specificity, we injected AstroHTP into the dorsal striatum (dSTR) of 8-week-old C57BL/6J mice and analyzed tissue 3 weeks later. We quantified viral expression using ^NLS^dTomato and determined cell-type identity by co-staining with nuclear markers of astrocytes (SOX9) (*16*), neurons (NeuN), and oligodendrocyte-lineage cells (OLIG2). AstroHTP labeled 53.2% ± 8.9% of dorsal striatal astrocytes (Fig. 1C–D). We did not detect dTomato expression in NeuN or OLIG2-positive nuclei. Thus, AstroHTP drove efficient, highly astrocyte-specific expression of HTP in striatal astrocytes.

We next asked whether HTP-expressing astrocytes effectively capture DART ligands *in vivo*. We infused the HTL-linked tracer dye, Alexa647^DART^, into the dSTR via an indwelling cannula and performed histology 24 hours later. Alexa647^DART^ accumulated robustly on ^NLS^dTomato-positive astrocytes, indicating efficient target engagement *in vivo* (Fig. 1E).

We next asked how long AstroHTP persists at the cell surface by measuring its turnover *in vivo* using a pulse-chase strategy. Biotin^DART^ was infused into the dSTR, followed by Alexa647^DART^ at intervals ranging from 1 to 72 hours. The remaining surface Biotin^DART^ was detected with streptavidin-Alexa750 (SA750) (Fig. 1F) and plotted as a function of chase interval (Fig. 1G). The decay was best fit by a double exponential (effective half-life: 1.41 ± 0.43 days), with a minor fast component accounting for the remaining 16% (half-life: 0.03 days) and a slow component accounting for 84% of the signal (half-life: 1.88 days). Thus, in astrocytes, most surface HTP persists for days, with a smaller, rapidly internalizing pool.

To confirm that astrocytes capture DART ligands onto Perisynaptic Astrocyte Processes (PAPs) at high resolution, we used a sparse transduction strategy, co-injecting a low-titer astrocyte-specific Cre virus with a high-titer Cre-dependent HTP-GPI-IRES-dTomato-farnesyl construct, in which farnesylated dTomato labels the inner membrane leaflet to enable reconstruction of the astrocyte’s surface. Following striatal Alexa647^DART^ infusion, the tracer signal colocalized robustly with dTomato-labeled astrocytic membranes (Fig. 1H), confirming ligand capture and establishing DART labeling as a proxy for the ramified architecture of astrocytic processes.

To further confirm the localization of AstroHTP to PAPs at ultrastructural resolution, we performed immuno-electron microscopy (immuno-EM). HTP-expressing striatal astrocytes were labeled *in vivo* with Biotin^DART^ (Fig.1I). Brains were fixed four hours later, and sections were incubated with Alexa Fluor 647-FluoroNanogold™-streptavidin to label Biotin^DART^-positive astrocytes. Regions of interest containing Alexa647 fluorescence were selected and prepared for EM (Fig. 1J, fig. S1A). EM revealed that nanogold particles were concentrated in PAPs and excluded from neuronal pre– and postsynaptic compartments (Fig. 1K, fig. S1B). Some of the AstroHTP-bound nanogold particles were detected intracellularly, consistent with internalization.

Finally, we tested whether AstroHTP could support DART ligand capture in other brain regions. AstroHTP was injected into the hippocampus or motor cortex, followed three weeks later by Alexa647^DART^ infusion. In both regions, Alexa647^DART^ robustly accumulated on ^NLS^dTomato-positive astrocytes, with no apparent labeling in neurons or oligodendrocyte-lineage cells (fig. S1C-E). Together, these results establish AstroHTP as an effective, astrocyte-specific DART platform that can be generalized across brain regions.

### Astrocyte-specific inhibition of AMPAR activity using YM90K^DART^

Transcriptomic and proteomic profiling studies show that striatal astrocytes express AMPAR subunits (*17, 18*) (Fig. S2A–B). Yet the functions of astrocytic AMPARs have remained difficult to test directly. AstroHTP provides a direct way to interrogate astrocytic AMPARs when paired with YM90K.1^DART.2^ (hereafter, YM90K^DART^), a tetherable competitive AMPAR antagonist (*11, 12*). Previous studies in HTP-positive neurons showed that low-dose exposure achieves potent AMPAR inhibition, because the concentration needed to drive drug tethering is approximately 300-fold lower than the reagent’s 50% ambient concentration (*AC*_50_) of 6 µM (fig. S2C). Once tethered, YM90K^DART^ is highly selective for AMPARs, with minimal off-target activity across other glutamate and non-glutamate receptors (*12*).

We tested whether YM90K^DART^ inhibits astrocytic AMPAR function in acute slices by using astrocytic calcium imaging as the readout (*8, 9*). 8-week-old C57BL/6J mice were injected with AstroHTP together with GfaABC1D-GCaMP6f, a fluorescent sensor of the cytosolic calcium concentration (*19*). AMPA (100 µM) was bath applied together with a positive allosteric modulator, cyclothiazide (CTZ, 100 µM), in the presence of tetrodotoxin (TTX, 0.5 µM) to inhibit neuronal action potentials. AMPA treatment under these conditions elicited robust astrocytic Ca²⁺ transients in striatal astrocytes, similar to those previously observed in cortical astrocytes (*20*). These responses were abolished by preincubation of slices with the AMPAR antagonist NBQX (30 µM), confirming that they required AMPAR activation (fig. S2D-F, Supp Video1-2). To determine if YM90K^DART^ can block AMPA-elicited astrocytic Ca²⁺ transients, slices were randomly assigned to preincubation with either 1 µM YM90K^DART^ or 1 µM Blank^DART^, a pharmacologically inert reagent (Fig. 2A-D). Both conditions also included 0.1 µM Alexa647^DART^, a fluorescent tracer that verifies ligand capture. Excess untethered ligand was washed out before imaging, ensuring that any effects of YM90K^DART^ reflect tethered, rather than ambient compound. Pre-incubation with YM90K^DART^ markedly reduced both the peak amplitude and the integrated Ca²⁺ signal in astrocytes relative to Blank^DART^ controls, confirming effective AMPAR antagonism (Fig. 2 E-G, fig. S2G, Supp Video3-4). We next examined the impact of YM90K^DART^-mediated astrocytic AMPAR inhibition on spontaneous astrocytic Ca^2+^ transients *ex vivo*. Striatal slices were again pre-treated with either YM90K^DART^ or Blank^DART^ but imaged in the absence of TTX, AMPA, or CTZ. Ca²⁺ events were analyzed with Activity Quantification and Analysis 2 (AQuA2), a machine-learning–based pipeline for resolving complex spatiotemporal calcium signals (*21*). YM90K^DART^ significantly reduced the frequency of spontaneous astrocytic Ca²⁺ events compared to slices treated with Blank^DART^, while mean event amplitude was unchanged (Fig. 2H-J, Supp Video 5-6). Thus, AstroHTP combined with YM90K^DART^ selectively inhibits astrocytic AMPAR signaling in striatal circuits.

**Figure 2.**
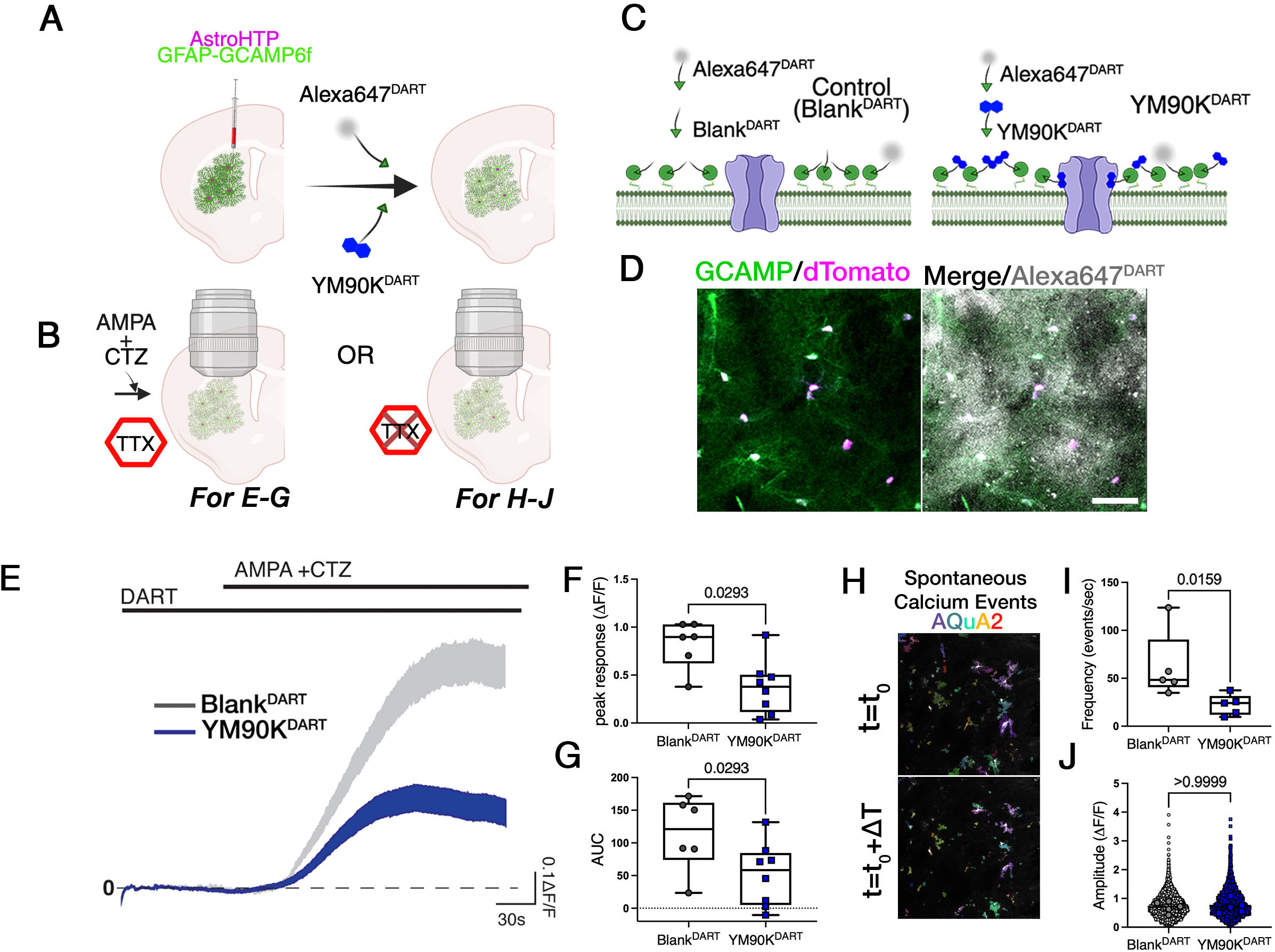
Inhibition of astrocytic AMPARs via YM90K^DART^. **(A)** Experimental schematic for inhibiting astrocyte AMPARs. Coronal striatal sections co-expressing AstroHTP and GFAP-GCaMP6f were pre-incubated with DART ligands. **(B)** Experimental schematic comparing two conditions: AMPA-evoked Ca²⁺ responses (100 µM AMPA + 100 µM cyclothiazide [CTZ] + 0.5 µM TTX; panels E–G) versus spontaneous Ca²⁺ signaling (AMPA omitted, TTX excluded; panels H–J). **(C)** Sections were incubated with either 1 µM Blank ^DART^ or 1 µM YM90K^DART^, together with 0.1 µM Alexa647^DART^ to visualize drug capture**. (D)** Representative image showing AstroHTP (dTomato, magenta) co-expressed with GCaMP6f (green) and Alexa647^DART^ capture (grey). Scale bar, 50 µm. **(E-G)** YM90K^DART^ attenuates AMPA-evoked Ca²⁺ responses in AstroHTP astrocytes. **(E)** ΔF/F traces showing Ca²⁺ responses of striatal astrocytes to bath application of 100 µM AMPA in the presence of 100 µM CTZ and 0.5 µM TTX. Traces correspond to slices pre-incubated with 1 µM Blank^DART^ (control, grey) or 1 µM YM90K^DART^ (blue). Shading represents mean ± SEM. **(F**) Summary of peak AMPA-evoked Ca²⁺ responses (ΔF/F): Blank^DART^, 0.822 ± 0.101; YM90K^DART^, 0.373 ± 0.0998 (Mann– Whitney test, P = 0.0293; n = 6 slices from 4 mice (Blank^DART^) and 8 slices from 5 mice (YM90KDART); 96 and 127 ROIs, respectively**). (G)** Summary of AMPA-evoked Ca²⁺ response AUC: Blank^DART^, 114 ± 22.9; YM90K^DART^, 51.8 ± 17.1 (Mann–Whitney test, P = 0.0293; n = 6 slices from 4 mice (Blank^DART^) and 8 slices from 5 mice (YM90K^DART^); 90 and 127 ROIs, respectively). Peak and AUC were analyzed using the same rank-based test; because these measures are highly correlated across slices, they yield identical U-statistics and P-values. **(H–J)** Astrocytic AMPARs regulate spontaneous Ca²⁺ signaling. **(H)** Representative images of astrocytic Ca²⁺ events detected using AQuA2. **(I)** The frequency of astrocytic Ca²⁺ events was reduced following YM90K^DART^ treatment (Blank^DART^ 62 ± 16; YM90K^DART^ 22 ± 4.8 events/s; Mann– Whitney test, P = 0.0159; n = 5 slices from 4 mice per condition). **(J)** Amplitude of astrocytic Ca²⁺ events was unchanged following YM90K^DART^ treatment (control 0.703 ± 0.0846; YM90K^DART^ 0.715 ± 0.104; Mann–Whitney test, P > 0.9999; n = 5 slices from 4 mice per condition).

### Unilateral inhibition of astrocytic AMPARs induces an ipsiversive turning bias

To determine whether astrocytic AMPAR signaling contributes to striatal-dependent behavior *in vivo*, we asked whether attenuating astrocytic AMPAR signaling alters locomotion in the open field. To maximize behavioral sensitivity, we focused on a unilateral perturbation with turning bias as the primary behavioral outcome. We injected AstroHTP into the left dSTR and implanted a cannula at the same location for intracranial ligand infusion. Three weeks after surgery, mice received direct striatal infusions of either 30 µM YM90K^DART^ or a control cocktail containing 30 µM untetherable YM90K^PEG36^ and 30 µM Blank^DART^. In both conditions, 3 µM Alexa647^DART^ was included to assess target engagement. Critically, because both conditions have the same quantity of PEG-linked YM90K, any ambient effects are matched across conditions. The HTP tethering process is likewise matched because equivalent amounts of PEG-linked HTL were used, leaving the presence or absence of a physical linkage between YM90K and HTL as the key experimental variable (Fig. 3A). To assess motor output, we recorded freely moving mice from both groups in an open-field arena for two 30-minute sessions, at 1 hour and 24 hours post-infusion, tracking 18 body parts using 3-D Aligned Neural Network for Computational Ethology (DANNCE) (*22, 23*) (Supp Video 7). YM90K^DART^-treated mice developed a directional bias 1 h post-infusion in the first session. This bias reflects two overlapping effects: cannula implantation and virus expression caused a contraversive (rightward) turning bias in controls (Fig. 3B), and YM90K^DART^ counteracted this bias by promoting turning in the ipsiversive (leftward) direction (Fig. 3B-D). This shift occurred despite no change in total turns (Fig. 3E) or total distance traveled (Fig.3F). By 24 hours post-infusion, the turning bias had largely resolved, with a trending but nonsignificant difference between groups (fig. S3A-D), suggesting that compensatory mechanisms may rebalance the circuit over time, underscoring the importance of an acute perturbation. Other secondary metrics, including distance traveled, time spent in the center, and time spent immobile, showed no effect of YM90K^DART^ relative to controls in either open-field session (fig. S3E-G). Given that contraversive movements require activity across direct– and indirect-pathway spiny projection neurons (*24*), the observed reduction in contraversive turns induced by astrocytic YM90K^DART^ suggests a net reduction in SPN activity.

**Figure 3.**
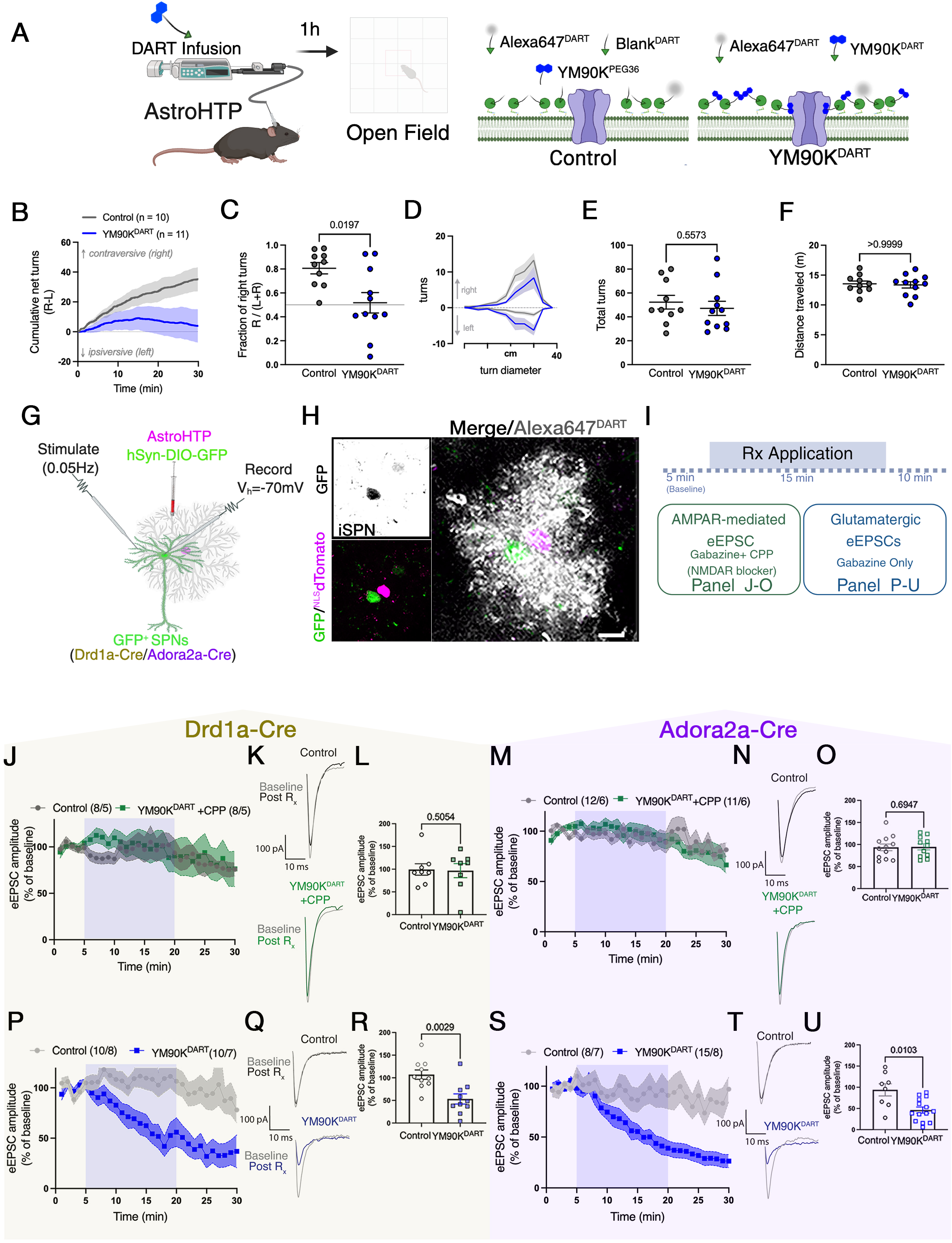
Astrocyte-specific inhibition of AMPARs reduces excitatory synaptic activity of SPNs in an activity-dependent and NMDAR-dependent manner. **(A)** Astrocytic AMPAR inhibition *in vivo*. Experimental schematic: adult C57BL/6J mice were injected with AstroHTP in the left dSTR and implanted with a cannula for intracranial DART infusion. Three weeks later, DART ligands were infused into head-fixed mice for 15 min, followed by open-field behavioral analysis. Mice were infused with 30 µM Blank^DART^ and 30 µM YM90K^PEG36^, or 30 µM YM90K^DART^, each together with 3 µM Alexa647^DART^. **(B-F)** Astrocytic AMPAR inhibition corrects cannula implantation contraversive (right) turning bias. **(B)** cumulative net turns of the mouse, showing both right and left turns over time and their accumulation over the session. Data are mean ± SEM, N = 10 control, 11 YM90K^DART^. **(C)** Fraction of right turns, treatment (Control 0.807± 0.0474; YM90K^DART^ 0.518± 0.0852; Mann– Whitney test, P = 0.0197; N = 10 control, 11 YM90K^DART^). **(D)** Turn-diameter histogram (1 cm bins) 1 h after control (grey) or YM90K^DART^ infusion. Left turns are plotted as downward bars and right turns as upward bars. Data are mean ± SEM, N = 10 control, 11 YM90K^DART^. **(E)** Total turns did not differ between groups (Control 52.3± 5.77; YM90K^DART^ 47.2± 5.92; Mann–Whitney test, P = 0.5573; N = 10 control, 11 YM90K^DART^). **(F)** Distance traveled did not differ between groups (Control 13.5± 0.475; YM90K^DART^ 13.4± 0.530; Mann–Whitney test, P >0.9999; N = 10 control, 11 YM90K^DART^). **(G)** Schematic of the viral strategy used to label SPNs and astrocytes. Drd1a-Cre and Adora2a-Cre mice were transduced with AAV5-hSyn-DIO-GFP to label direct-pathway SPNs (dSPNs) and indirect-pathway SPNs (iSPNs), respectively, together with AstroHTP, which expresses HaloTag protein (HTP) and NLS-dTomato in astrocytes. GFP-positive SPNs located near NLS-dTomato-positive astrocytes were patch-clamped (held at −70 mV) to measure evoked excitatory postsynaptic currents (eEPSCs). **(H)** Representative image of a GFP-positive iSPN adjacent to an NLS-dTomato-positive, HTP-expressing astrocyte capturing Alexa647^DART^ (grey). Scale bar, 10 µm. **(I)** Slice pharmacology conditions and recording paradigms for panels J-O and P-U. *Pharmacology:* control, 300 nM Blank^DART^ + 300 nM YM90K^PEG36^ + 30 nM Alexa647^DART^; YM90K^DART^, 300 nM YM90K^DART^ + 30 nM Alexa647^DART^. *AMPAR-mediated eEPSCs (J–O):* recorded in the presence of gabazine and CPP to block GABA_A_ receptor– and NMDA receptor-mediated currents, respectively, isolating the AMPAR component of the eEPSC. *Glutamatergic eEPSCs (P-U):* recorded in gabazine alone, with CPP omitted, so NMDA receptor-mediated currents remain intact alongside the AMPAR component. **(J–O)** Astrocyte-tethered YM90K^DART^ does not act transcellularly on neuronal AMPARs. **(J)** Time course of normalized eEPSC amplitude in dSPNs. In the presence of CPP, YM90K^DART^-treated cells (green) showed no change relative to control (grey). Data are mean ± SEM. **(K)** Representative eEPSC traces from dSPNs at baseline (grey) and after treatment with control (black) or YM90K^DART^ (green). **(L)** End-point comparison of dSPN eEPSC amplitude (CPP condition): control 99.3 ± 12.6%; YM90K^DART^ 96.8 ± 15.5% (Mann–Whitney test, P = 0.5054). Control, 8 cells, 5 mice; YM90K^DART^, n = 8 cells, 5 mice. **(M)** Time course of normalized eEPSC amplitude in iSPNs. In the presence of CPP, YM90K^DART^ treated cells (green) showed no change relative to control (grey). Data are mean ± SEM. **(N)** Representative eEPSC traces from iSPNs at baseline (grey) and after treatment with control (black) or YM90K^DART^ (green). **(O)** End-point comparison of iSPN eEPSC amplitude: Control 93.5 ± 7.63%; YM90K^DART^ 94.6 ± 6.94% (Mann–Whitney test, P = 0.6947). Control, n = 12 cells, 6 mice; YM90K^DART^, n = 11 cells, 6 mice. (**P-U)** Astrocyte-specific inhibition of AMPARs reduces excitatory synaptic activity of SPNs. **(P)** Time course of normalized dSPN eEPSC amplitude; YM90K^DART^ (blue) showed a progressive reduction compared with control (grey). Data are mean ± SEM. **(Q)** Representative eEPSC traces from dSPNs at baseline (grey) and after treatment with control (black) or YM90K^DART^ (blue). **(R)** End-point comparison of dSPN eEPSC amplitude: control 108 ± 8.6%; YM90K^DART^ 62 ± 9.59% (Mann–Whitney test, P = 0.0029). Control, n = 10 cells, 8 mice, YM90K^DART^, n = 10 cells, 7 mice. **(S)** Time course of normalized eEPSC amplitude in iSPNs; YM90K^DART^ (blue) showed a progressive reduction compared with control (grey). Data are mean ± SEM.**(T)** Representative eEPSC traces from iSPNs at baseline (grey) and after treatment with control (black) or YM90K^DART^ (blue). **(U)** End-point comparison of iSPN eEPSC amplitude: control 93.9 ± 14.4%; YM90K^DART^ 48.0 ± 6.93% (Mann–Whitney test, P = 0.0103). Control n = 8 cells, 7 mice, YM90K^DART^ n = 15 cells, 8 mice.

### Astrocyte-tethered YM90K^DART^ does not act transcellularly on neuronal AMPARs

We next asked whether an astrocyte-tethered YM90K^DART^ could act transcellularly to inhibit neuronal AMPARs. Drd1a-Cre or Adora2a-Cre mice were injected with AAV-hSyn-DIO-GFP to label genetically identified direct– or indirect-pathway SPNs (dSPN or iSPN, respectively), while striatal astrocytes were transduced with AstroHTP. Three to four weeks later, we prepared acute coronal brain slices and obtained whole-cell voltage-clamp recordings from GFP-positive SPNs adjacent to ^NLS^dTomato^+^ astrocytes (Fig. 3G-H). We isolated neuronal AMPAR-mediated currents by pharmacologically blocking GABA_A_ and NMDA receptors. Evoked EPSCs (eEPSCs) were elicited every 20 seconds to monitor AMPA-mediated synaptic responses. After a stable 5-minute baseline, we bath-applied YM90K^DART^ or the control cocktail for 15 minutes, followed by a 10-minute washout during which recordings continued (Fig. 3I). Under these conditions, YM90K^DART^ did not alter eEPSC amplitude in either dSPNs (Fig. 3J-L) or iSPNs (Fig. 3M-O). By contrast, when HTP was expressed in SPNs, the neuronally-tethered YM90K^DART^ robustly reduced AMPAR-mediated eEPSC amplitude (fig. S4). Thus, astrocyte-tethered YM90K^DART^ does not appreciably cross the perisynaptic cleft to engage neuronal AMPARs, consistent with previously shown membrane-confined action (*11, 12, 25*).

### Astrocytic glutamate sensing through AMPARs is necessary to maintain synaptic strength

We next repeated the experiment in the absence of NMDAR antagonism. Under these conditions, inhibition of astrocytic AMPARs via YM90K^DART^ produced a robust reduction in eEPSC amplitude in both dSPNs (Fig. 3P-R) and iSPNs (Fig. 3S-U). This suppression persisted after washout of free ligand, further confirming that the effect arises from stable tethering of YM90K^DART^ to AstroHTP rather than from residual ambient compound. These results suggest that astrocytic AMPAR inhibition induces a robust, NMDAR-dependent decrease in synaptic strength that resembles long-term depression (LTD) (*26*).

To determine whether this decrease in synaptic strength requires ongoing synaptic transmission (*27, 28*), we compared continuous afferent stimulation throughout YM90K^DART^ application to a protocol in which afferent stimulation was paused during YM90K^DART^ application and resumed following washout (Fig. 4A). As before, YM90K^DART^ treatment reduced eEPSC amplitude when paired with ongoing synaptic stimulation (Fig. 4B-D). By contrast, when stimulation was paused during YM90K^DART^ application, the first post-drug eEPSC was unchanged from baseline (Fig. 4B-C, E-F), and eEPSC depression reemerged following the resumption of stimulation (Fig. 4G-H, fig. S6A). Similarly, astrocytic AMPAR inhibition had no effect on the amplitude or frequency of miniature excitatory postsynaptic currents (mEPSCs) in the absence of NMDAR antagonism without evoked stimulation (fig. S5). Taken together, these results show that blocking astrocytic AMPARs reduces synaptic strength in an activity-dependent manner, consistent with a use-dependent LTD-like phenomenon.

**Figure 4.**
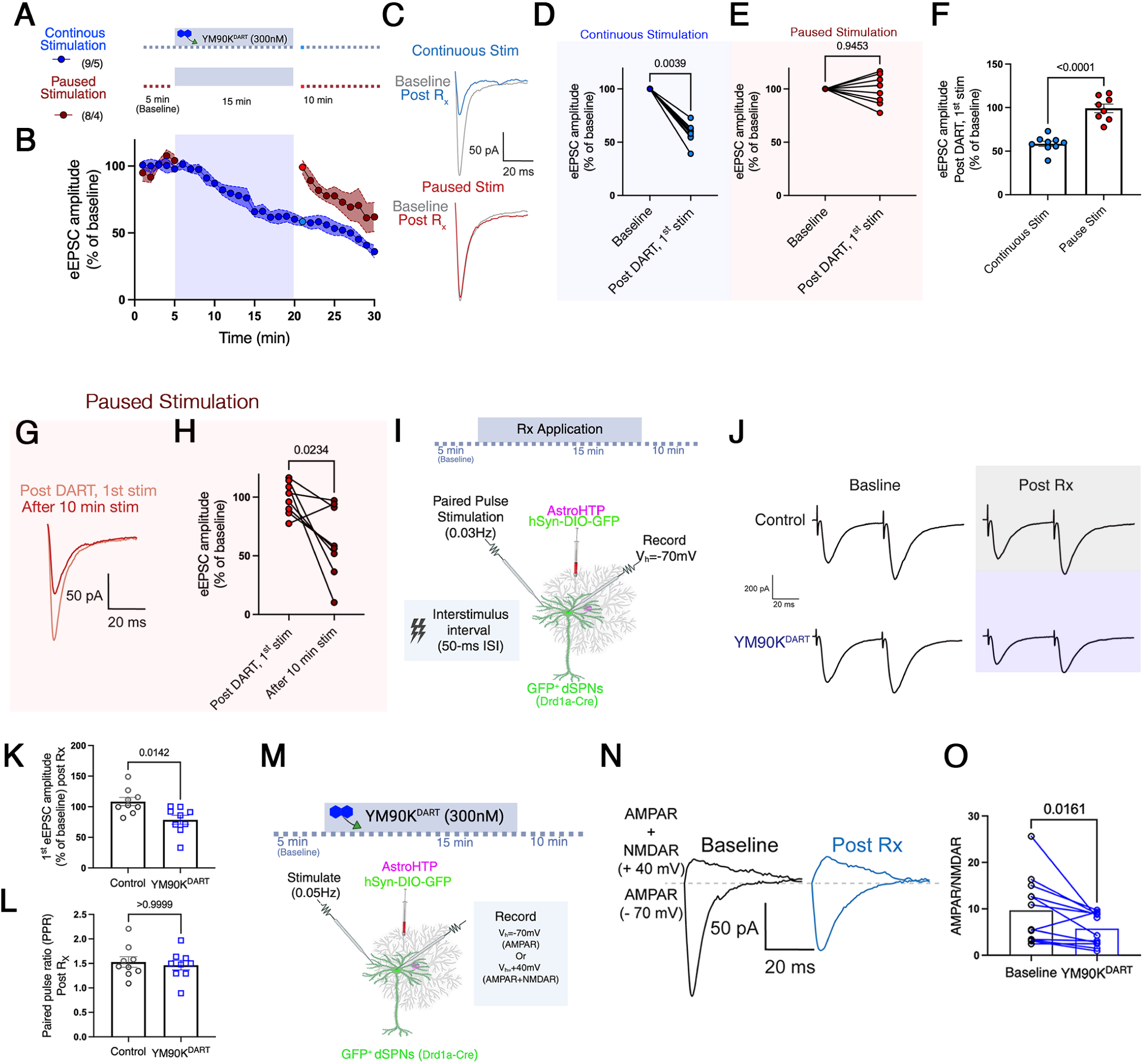
Astrocyte-specific AMPAR inhibition induces use-dependent postsynaptic LTD.(A-H) Astrocyte-specific inhibition of AMPARs reduces excitatory synaptic activity in dSPNs in an activity-dependent manner. **(A)** Recording paradigm for panels B-F. eEPSCs were recorded as described in Figure 3. Both groups received YM90K^DART^ (300 nM). In the continuous-stimulation group, synaptic stimulation continued throughout the baseline period, YM90K^DART^ application, and washout. In the paused-stimulation group, stimulation was suspended at the end of a 5-min baseline period, remained off during YM90K^DART^ application, and resumed after 15 min of drug exposure. **(B)** Time course of normalized eEPSC amplitude in dSPNs. The continuous-stimulation group (blue) exhibited a robust reduction in eEPSC amplitude during continuous synaptic stimulation. In contrast, the paused-stimulation group (red) showed stable eEPSC amplitudes immediately following resumption of stimulation after YM90K^DART^ exposure, followed by a progressive decrease with continued synaptic stimulation. Data are mean ± SEM. Continuous stimulation, n = 9 cells, 5 mice; paused stimulation, n = 8 cells, 4 mice**. (C)** Representative eEPSC traces from dSPNs at baseline (grey) and after YM90K^DART^ treatment with continuous stimulation (blue) or paused stimulation (red). **(D)** Paired comparison of eEPSC amplitude at baseline and after 15 min YM90K^DART^ application with continuous stimulation, 58.5 ± 3.01% (Wilcoxon test, P = 0.0039), n=9 cells, 5 mice. **(E)** Paired comparison of eEPSC amplitude at baseline and after 15 min YM90K^DART^ application with paused stimulation, 99.1 ± 4.93% (Wilcoxon test, P = 0.9453), n=8 cells, 4 mice. **(F)** Comparison of the first eEPSC amplitude after 15 min of drug application between the continuous– and paused-stimulation groups: continuous 58.5 ± 3.01%; paused 99.1 ± 4.93% (Mann–Whitney test, P < 0.0001). Continuous stimulation, n = 9 cells, 5 mice; paused stimulation, n = 8 cells, 4 mice**. (G)** Representative eEPSC traces from a dSPN during the paused-stimulation protocol, first after stimulation resumed (light red) and 10 minutes after continuous stimulation resumed (dark red). **(H)** Paired comparison of eEPSC amplitude before versus after the stimulation pause: first stimulation, 99.1 ± 4.93%; after 10 min of resumed stimulation, 62.0 ± 10.9% (Wilcoxon test, P = 0.0234), n = 8 cells, 4 mice**. (I-L)** Astrocyte-specific inhibition of AMPARs does not change presynaptic release probability. **(I)** Stimulation and recording paradigm for paired-pulse ratio (PPR) measurements, applied throughout panels J-K. Paired-pulse stimuli were delivered at 0.03 Hz, with a 20 Hz (50 ms) interstimulus interval. Pharmacology; control: 300 nM Blank^DART^ + 300 nM YM90K^PEG36^ + 30 nM Alexa647^DART^; YM90K^DART^: 300 nM YM90K^DART^ + 30 nM Alexa647^DART^. **(J)** Representative PPR traces in control and YM90K^DART^-treated conditions. Control traces are shown at baseline and post-treatment (black); YM90K^DART^-treated traces are shown at baseline and post-treatment (blue). **(K)** Quantification of the first eEPSC amplitude in the PPR experiment, normalized to baseline. YM90K^DART^ treatment reduced eEPSC amplitude (control 108 ± 6.77% of baseline YM90K^DART^ 78.6 ± 7.25% of baseline; Mann–Whitney test, P = 0.0142). **(L)** Quantification of PPR, showing no change following YM90K^DART^ treatment (control 1.53 ± 0.109; YM90K^DART^ 1.46 ± 0.0982; Mann–Whitney test, P > 0.9999). Both measures: n = 9 cells from 5 mice. **(M-O)** Astrocyte-specific inhibition of AMPARs reduces postsynaptic AMPAR signaling. **(M)** Recording paradigm for AMPAR/NMDAR ratio measurements characterized in N-O. **(N)** Representative AMPAR/NMDAR traces recorded at −70 mV (AMPAR) and +40 mV (NMDAR) in dSPNs, at baseline (black) and post-YM90K^DART^ (blue). **(O)** Quantification of AMPAR/NMDAR ratio at baseline and after YM90K^DART^ treatment: baseline 9.69 ± 2.05; post-Rx 5.74 ± 1.01 (Wilcoxon matched-pairs signed-rank test, P = 0.0161; n = 12 cells, 5 mice).

In the striatum, LTD is canonically expressed presynaptically, via a decrease in glutamate release probability (*29*), and a resulting increase in the paired pulse ratio (PPR; second divided by first eEPSC amplitude). We thus repeated the assay in dSPNs adjacent to AstroHTP-expressing astrocytes while measuring PPRs (Fig. 4I-J). As before, astrocytic YM90K^DART^ induced a marked reduction in eEPSC amplitude (Fig. 4K, fig. S6B). Surprisingly, in contrast to canonical LTD in these cells, the PPR remained similar between control and YM90K^DART^-treated groups (Fig. 4L, fig. S6C). We then examined postsynaptic glutamate receptor function (Fig. 4M). We found that astrocytic AMPAR inhibition, combined with evoked stimulation in the absence of NMDAR blockers, significantly reduced the AMPAR/NMDAR ratio (Fig. 4N-O). Taken together, these findings indicate that the observed LTD is primarily driven by a reduction in postsynaptic AMPAR-mediated currents.

### Astrocyte-tethered YM90KDART induces LTD via non-ionotropic NMDAR signaling

Although postsynaptic LTD is not typical for SPNs, it is the canonical form of LTD for pyramidal neurons, wherein NMDAR-dependent LTD classically requires Ca²⁺ influx, activating phosphatases that drive AMPAR internalization and a resulting drop in the AMPAR/NMDAR ratio (*30, 31*). However, NMDARs can also induce LTD non-ionotropically, wherein glutamate induces a conformational change that triggers intracellular signaling independent of ion flux (*32*).

Even though Mg²⁺ blocks ionotropic NMDAR signaling at our −70 mV holding potential (*33*), ionotropic signaling may still occur in spines or in dendritic compartments that are not fully space-clamped (*34*). Thus, to decisively assess whether ionotropic NMDAR signaling is required, we co-applied the NMDAR pore blocker MK-801 with YM90K^DART^ (Fig. 5A). MK-801 did not prevent the reduction in eEPSC amplitude induced by YM90K^DART^ (Fig. 5B-D, fig. S6D-E), indicating that ionotropic NMDAR signaling is not required for this form of LTD.

**Figure 5.**
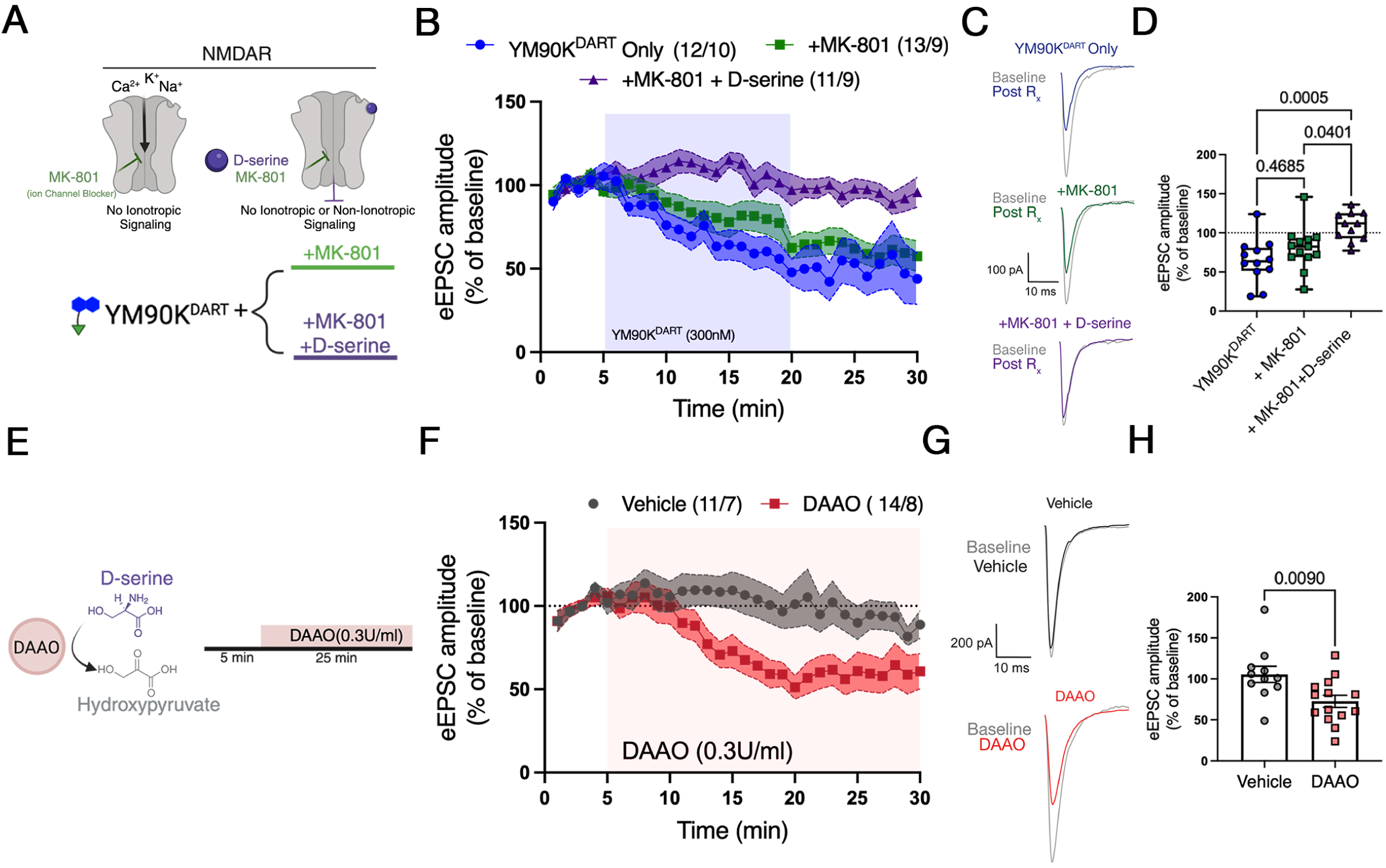
Astrocyte-specific inhibition of AMPARs reduces excitatory synaptic activity via non-ionotropic NMDAR signaling. **(A)** Schematic of the eEPSC recording paradigm used in panels B-D. Whole-cell dSPN recordings were performed at −70 mV in the presence of gabazine alone, gabazine plus MK-801 (to block ionotropic NMDAR activity), or gabazine plus MK-801 and D-serine (to inhibit both ionotropic and non-ionotropic NMDAR signaling). Astrocytic AMPARs were inhibited by bath application of YM90K^DART^ (300 nM). **(B)** Time course of normalized eEPSC peak amplitude in dSPNs. YM90K^DART^ alone (blue) induced a progressive reduction in eEPSC amplitude that persisted in the presence of MK-801 (green) but was abolished by co-application of MK-801 and D-serine (purple). Data are mean ± SEM (YM90K^DART^, 12 cells from 10 mice; YM90K^DART^ + MK-801, 13 cells from 9 mice; YM90K^DART^ + MK-801 + D-serine, 11 cells from 9 mice). **(C)** Representative eEPSC traces recorded at baseline (grey) and after YM90K^DART^ alone (blue), YM90K^DART^ + MK-801 (green), or YM90K^DART^ + MK-801 + D-serine (purple). **(D)** End-point eEPSC amplitude (10– 20 min post-infusion) differed across conditions (Kruskal–Wallis test, P = 0.0007). Dunn’s post hoc comparisons revealed no difference between YM90K^DART^ and YM90K^DART^ + MK-801 (P = 0.4685), but significant differences between YM90K^DART^ and YM90K^DART^ + MK-801 + D-serine (P = 0.0005), and between YM90K^DART^ + MK-801 and YM90K^DART^ + MK-801 + D-serine (P = 0.0401). **(E-H)** D-serine depletion reduces excitatory synaptic activity in dSPNs. **(E)** Schematic of the eEPSC recording paradigm used in panels F-H. Recordings were performed at −70 mV in the presence of gabazine. D-serine depletion was achieved by bath application of D-amino acid oxidase (DAAO, 0.3 U/ml). **(F)** Time course of eEPSC amplitude in dSPNs showing that DAAO (red) reduced eEPSC amplitude relative to vehicle (grey). Data are mean ± SEM (vehicle, 11 cells from 7 mice; DAAO, 14 cells from 8 mice). **(G)** Representative eEPSC traces at baseline (grey) and after vehicle (black) or DAAO (red). **(H)** End-point eEPSC amplitude (10–20 min post-infusion) was reduced in DAAO-treated slices compared with vehicle (vehicle 105 ± 9.96%; DAAO 72.7 ± 7.40%; Mann–Whitney test, P = 0.0090).

Non-ionotropic NMDAR signaling (*35, 36*) is strongly attenuated by NMDAR co-agonists (*37*), and D-serine is the main endogenous NMDAR co-agonist in the striatum (*38, 39*). We thus combined MK-801 with a saturating concentration of D-serine (10 µM) to suppress non-ionotropic NMDAR signaling. Under these conditions, astrocyte-tethered YM90K^DART^ no longer induced LTD (Fig. 5B-D, fig. S6F). These results indicate that astrocytic AMPAR inhibition mediates LTD via non-ionotropic NMDAR signaling and suggest that NMDAR co-agonists, such as D-Serine, are critical for maintaining synaptic strength during ongoing synaptic activity.

Finally, we tested whether NMDAR co-agonist depletion is sufficient to phenocopy astrocytic AMPAR inhibition. To do so, we bath-applied D-amino acid oxidase (*40*) (DAAO; 0.3 U/mL), an enzyme that degrades D-serine, glycine, and other NMDAR co-agonists, whilst using the same eEPSC recording paradigm in Fig. 3G. Importantly, we did not use YM90K^DART^ in these experiments. Under these conditions, DAAO alone reduced eEPSC amplitude to a similar extent as astrocytic YM90K^DART^ (Fig. 5E-H, fig. S6G-H), indicating that reduced NMDAR co-agonist availability is sufficient to recapitulate the effect of astrocytic YM90K^DART^. Together, these results show that an extracellular NMDAR co-agonist, such as D-serine, is required for preserving synaptic strength during ongoing synaptic activity. Taken together with previous results, these findings also suggest that astrocytic AMPARs maintain synaptic strength by regulating extracellular NMDAR co-agonist availability.

## Discussion

A growing body of work positions astrocytes as an instructive element that can shape circuit formation and function in the striatum (*40–43*). However, the functional consequences of neurotransmitter sensing by perisynaptic astrocytic receptors remain poorly understood. Our results reveal a fundamental role for AMPAR-mediated astrocytic glutamate sensing as a parity signal required to preserve synaptic strength during use. In the absence of this parity signal, non-ionotropic NMDAR signaling induces a parity-mismatch form of LTD.

In the striatum, learning-related LTD is canonically instructed by spike timing and by dopamine, endocannabinoid, adenosine, and cholinergic signaling, and is expressed presynaptically (*44–52*). Parity-mismatch LTD is fundamentally different: it arises when glutamate transmission occurs without a matching astrocyte-dependent signal, and it is expressed postsynaptically. Whether it also contributes to learning-related plasticity remains unknown.

Our findings carry implications for striatal pathophysiology. Maladaptive changes in corticostriatal synaptic strength have been implicated in Parkinson’s disease, levodopa-induced dyskinesia, and obsessive-compulsive disorder, which have all been associated with astrocytic involvement (*18, 53–55*). Loss of the astrocytic AMPAR-dependent parity signal induces LTD onto both direct– and indirect-pathway SPNs, suggesting a mechanism by which astrocytic dysfunction could globally suppress striatal gain (*29, 50*). Consistent with this possibility, striatal D-serine dysregulation has been documented in Parkinson’s disease and levodopa-induced dyskinesia (*56*). More broadly, the astrocytic parity signal provides a framework whereby synaptic pathologies can arise from failed astrocytic function rather than a primary neuronal defect, a hypothesis that is now directly testable in disease models using AstroHTP.

Several mechanistic questions remain. Astrocytic AMPAR inhibition reduced the frequency of spontaneous astrocytic calcium events, but whether these dynamics contribute to parity-mismatch LTD is unresolved (*57–59*). It is also unclear if astrocytes regulate AMPAR trafficking within peripheral processes, analogous to neuronal activity-dependent trafficking (*60, 61*), and whether astrocytic AMPAR inhibition alters the astrocytic membrane potential in ways that contribute to parity-mismatch LTD (*62*). The identity of the parity signal also remains open: D-serine is thought to predominate at striatal synapses, but DAAO can also oxidize other NMDAR coagonists such as glycine, albeit with a Km roughly 1000-fold above physiological glycine concentrations (*63*), so our enzymatic approach cannot fully distinguish the contributions of D-serine, glycine, or other D-amino acids to preserving synaptic strength downstream of astrocytic AMPAR engagement. Finally, why astrocytes, rather than neurons, provide the parity signal is a major question. Because each astrocyte ensheathes many synapses (*64, 65*), an astrocyte-derived parity signal may operate at a spatial scale larger than that of an individual synapse, positioning astrocytes to integrate local circuit context into parity signaling decisions. Defining the spatiotemporal spread of this signal remains an important goal of future work.

In conclusion, these findings identify an astrocytic parity signal that stabilizes synaptic strength by linking glutamate sensing to suppression of non-ionotropic NMDAR signaling. Astrocyte-targeted DART was essential for revealing this mechanism because it enables acute, cell-specific inhibition of endogenous receptors in intact circuits. Extending this strategy across other astrocytic receptor classes and disease models should reveal how these glial cells decode neurotransmitter signals to shape neural computation.

## Acknowledgments

This work was supported by National Institutes of Health BRAIN Initiative Grants to CE. And G.Y. (U19NS123719) and to MRT (RF1-MH117055 and R01-MH132592) and by the joint efforts of MJFF and the Aligning Science Across Parkinson’s (ASAP) initiative. MJFF administers the ASAP-020607 grant to CE and MRT. NE was funded by a Ruth K. Broad Foundation and a Life Sciences Research Foundation Postdoctoral Fellowship. C.E. is an HHMI investigator.

## Author contributions

Conceptualization: NE, JRR, JK, MRT, CE

Methodology: NE, JRR, JK, TV, BCS, GY, MRT, CE

Investigation: NE, JRR, JK, BCS, OWF, KU, SSXL, JMR, WZ, XC, MEvP, HY

Visualization: NE, JRR, JK, TV, HY, MRT, CE

Funding acquisition: NE, GY, MRT, CE

Project administration: NE, GY, MRT, CE

Supervision: GY, MRT, CE

Writing – original draft: NE, JRR, JK, MRT, CE

Writing – review & editing: NE, JRR, JK, TV, HY, KU, GY, MRT, CE

## Data, code, and materials availability

All data and custom code are available at DOI: https://doi.org/10.70106/r4583. Further information and requests for resources, reagents, data, and code should be directed to and will be fulfilled by the corresponding author.

## Supplementary Materials

### Materials and Methodes

#### Animals

Mice aged 6–16 weeks were used for all experiments. Wild-type C57BL/6J mice, as well as transgenic Drd1a-cre [B6.FVB(Cg)-Tg(Drd1a-cre) EY262Gsat/Mmucd] and Adora2a-Cre [B6.FVB(Cg)-Tg(Adora2a-cre) KG139Gsat/Mmucd]; mouse lines were used to selectively target striatal direct– and indirect-pathway medium spiny neurons, respectively. Mice were housed under standard conditions, with a 12-h light/dark cycle and ad libitum access to food and water. For behavioral experiments, mice were housed on a reverse 12h/12h light-dark cycle, and experiments were conducted during the animals’ active phase between zeitgeber times (ZT) 14 and 20 under dim red-light conditions. All experimental procedures were approved by the Institutional Animal Care and Use Committee (IACUC) and conducted in accordance with NIH guidelines.

#### AAV Production

AAVs were produced as previously described (*66*). In brief, HEK 293T cells were transfected with pAd-ΔF6, serotype plasmid AAV PHP.AX, and AAV transfer plasmid. After 72 h, the cells were lysed in 15 mM NaCl and 5 mM Tris-HCl (pH 8.5) and incubated with 50 U/ml benzonase for 30 min at 37 °C. The cell lysate was then centrifuged at 4,500 rpm for 30 min at 4 °C. The supernatant containing AAV was added to the top of an iodixanol gradient (15%, 25%, 40%, and 60% iodixanol solutions, from top to bottom) and centrifuged using a Beckman Ti-70 rotor at 60,000 rpm for 100 min. The viral solution was extracted from the virus layer (between the 40% and 60% iodixanol layers) using a 24-gauge needle and a 5-ml syringe, then concentrated using a 100-kDa filter. Viral titers were measured by quantitative PCR using a linearized plasmid containing the viral genome as the standard (Addgene). In addition, AAV5-GfaABC1D-GCaMP6f and AAV5-hSyn-DIO-GFP were purchased from Addgene Viral Service. Some batches of AAV-PHP.AX– GfaABC1D–HTP–NLS-dTomato–4×6T were custom-produced and purchased from VectorBuilder Inc.

#### Viral Surgery

Adult mice (6–16 weeks old) were anesthetized with 2% isoflurane and secured in a stereotaxic instrument (Model 1900, Kopf Instruments) on a heating pad set to 37 °C to maintain body temperature. Extended-release buprenorphine (0.5 mg/kg) was given subcutaneously and lubricating ocular ointment was applied. The scalp was shaved and then cleaned with alternating iodine and ethanol swabs. An incision was made along the midline of the skull, with 0.25% bupivacaine applied to the edges of the incision, then the surface of the skull was cleaned with 3% hydrogen peroxide. The skull was then leveled along the anterior/posterior and medial/lateral axes. Craniotomies were drilled at desired coordinates with an Alpen carbide drill bit (R1001/4c, Coltene, Inc.) using a Midwest Tradition TL drill (780344, Dentsply Sirona). Phosphate buffered saline (PBS) with 200 mM NaCl (to prevent aggregation) was used to dilute AAVs. Virus was loaded into a glass micropipette (Wiretrol II 5 µL, 5-000-2005, Drummond Scientific Company) and injected through pre-drilled craniotomies using a manual injector (MO-10, Narishige) at ∼100 nL/min. For dSTR targeting, virus was delivered across multiple injection sites to ensure uniform coverage (120 nL per depth). After each injection, the pipette was left in place for 3 to 5 min before retraction. Coordinates relative to bregma were AP +0.05, ML −1.7, DV −2.8 and −2.3; AP +0.15, ML −2.7, DV −3.2 and −2.7; AP +0.85, ML −1.3, DV −3.0 and −2.5; and AP +0.95, ML −2.3, DV −3.2 and −2.7. For motor cortex injections, 150 nL virus was delivered per site at AP +2.2, ML −0.9, DV −1.3; AP +2.2, ML −1.6, DV −1.6; AP +1.5, ML −0.8, DV −1.3; and AP +1.5, ML −1.6, DV −1.7. For hippocampus injections 500 nL virus was delivered at one site, AP −2.5, ML −1.7, DV −2.3. DV measurements are relative to skull surface at bregma. For surgeries not involving cannula implantation, the incision was closed with absorbable sutures (5-0 Webcryl, Pivetal), and mice were monitored during recovery and kept on a heating pad. Mice were used for experiments at least 3 weeks after viral surgery to allow for expression and recovery.

#### Cannula Implantation

For experiments involving intracranial DART delivery, guide cannulas were implanted immediately after viral injection. For striatal infusion, single guide cannulas (26 GA, guide cannula length 2 mm, dummy cannula 1 mm projection, Protech International or RWD Life Science with matched specifications) were implanted centered at the injection site (AP+0.5, ML −1.9, DV −2.5, [from skull surface at bregma, DV from the tip of the dummy cannula]). For intracerebroventricular access, bilateral cannulas (26 GA, center-to-center 2 mm, guide cannula length 2 mm, dummy cannula projection 0 mm, Protech International) were implanted targeting the lateral ventricles (AP −0.75, ML ±1.0, DV −2.25, [from skull surface at bregma, DV from the tip of the dummy cannula]). After slowly lowering cannulas, they were initially fixed in place using UV-curing Optibond (29669, Kerr Dental). A custom 3D-printed ‘headbar’ (*67*) was placed on the skull and secured in place with dental cement (Teets Cold Cure Denture Material powder and liquid, Coral, Co-Oral-Ite Dental Mfg.). For co-housed mice, an additional custom 3D-printed ‘headhat’ (*67*) was secured to the headbar to protect the cannula. Mice were used for experiments at least 3 weeks after viral surgery, allowing for expression and recovery.

#### Intracranial DART Delivery

DART infusions were performed in awake animals, head-fixed via the implanted headbar, using an automated syringe pump (PHD2000, Harvard Apparatus) connected to a 5 µL syringe (Model 75 RN, Hamilton) and a metal internal cannula (Protech International or RWD Life Science) extending 1.5 mm beyond the guide cannula for the striatum and 0.5 mm beyond the guide cannula for the lateral ventricle. Internal cannulas were connected to the syringe via PE50 tubing (0.023’’ ID × 0.050’’ OD, Protech International). Before infusion, the dummy cannula was removed, and the internal cannula was inserted. A total volume of 0.9 µL was infused at a rate of 0.1 µL/min, allowing an additional 6 min for diffusion before removing the internal cannula and replacing it with the dummy cannula. All DART reagents were prepared in Alzet ACSF. For infusions of YM90K^DART^ or YM90K^PEG36^, a plastic internal cannula (PEEK, Protech International) was used to avoid potential reductions in the nitroaromatic structure of YM90K caused by metal. For all other infusions, a metal internal cannula was used.

To visualize DART ligand capture *in vivo*, we infused Alexa Fluor 647 ^DART^ ^(^3µM, 0.9 µL total for 15 minutes) together with either control cocktail containing 30 µM untetherable YM90K^PEG36^ and 30 µM Blank^DART^ or 30µM YM90K^DART^ into the dSTR via an indwelling cannula and performed histology 24 hours later.

#### Histology and Immunostaining

Tissue processing for histology was performed using two protocols, reflecting the standard procedures of the laboratories where the analyses were conducted. The protocol for each experiment is indicated below.

##### Cryostat sections

Mice were anesthetized with tribromoethanol (200 mg/kg, Avertin) and transcardially perfused with TBS containing heparin (10 units/mL), followed by 4% paraformaldehyde (PFA) in TBS. Brains were dissected and post-fixed in 4% PFA overnight at 4 °C, cryoprotected in 30% sucrose, frozen in a 2:1 mixture of 30% sucrose and O.C.T. (TissueTek), and stored at −80 °C. Coronal sections (40–50 µm) were cut on a cryostat (CM3050S, Leica), collected as free-floating sections, and stored in a 1:1 mixture of TBS and glycerol at −20 °C until use.

##### Vibratome sections (Alexa647^DART^ capture)

Mice were deeply anesthetized by isoflurane inhalation in a closed chamber with physical separation from the anesthetic until cessation of breathing and loss of response to toe and tail pinch, then transcardially perfused with PBS followed by 4% PFA in PB. Brains were dissected, post-fixed in 4% PFA overnight, then washed in PBS. Brains were embedded in 4% agarose, then coronal sections (50 µm) were cut on a vibratome (VT1200S, Leica). Alternating sections were mounted onto glass slides or collected as free-floating sections and stored in PBS at 4 °C until use. For experiments visualizing native fluorescence only (e.g., Alexa647^DART^ capture), slides were air-dried, rehydrated in PBS, rinsed with water, then coverslipped with Vectashield Vibrance Antifade Mounting Medium with DAPI (Vector Labs) and allowed to cure overnight at room temperature. Slides were imaged on an Olympus FV3000 confocal microscope or an Olympus VS200 Slide Scanner with filter cubes for DAPI, TRITC, and Cy5. For experiments requiring antibody labeling, sections were processed as described under ‘Immunostaining’ below.

##### Immunostaining

The following protocol was applied to both free-floating cryostat and vibratome sections. Sections were blocked in PBS containing 0.1% Triton X-100 and 10% normal goat serum (NGS) for 1–2 h at room temperature, then incubated with primary antibodies (Sox9 1:2000; Olig2 1:500; NeuN 1:1000; see Key Resources Table for list) diluted in the same buffer for 1–2 days at 4 °C. Sections were then incubated with Alexa Fluor 488-, 568-, or 647-conjugated secondary antibodies (1/250, see Key Resources Table for list) diluted in the same buffer for 2 h at room temperature. Sections were mounted in Vectashield Vibrance Antifade Mounting Medium with Dapi (Vector Labs). Confocal images were acquired on an Olympus FV3000 or Leica SP8 microscope with filter cubes for DAPI, FITC, TRITC, and Cy5.

#### HaloTag Surface Turnover

##### Pulse-chase experiment

Adult mice (8 weeks old) were stereotaxically injected with AAV.PHP.AX-GfaABC1D-HTP-NLSdTomato-4×6T (2 × 10^12^ vg/mL) into the dSTR and implanted with a guide cannula, as described above. Three weeks post-surgery, mice were head-fixed and received a unilateral intracranial infusion of 0.9 µL of 60 µM Biotin^DART^. This pulse was followed, at intervals ranging from 1 to 72 h, by an infusion of 0.9 µL of 60 µM Alexa647^DART^. 24 h after the second infusion, brains were collected and processed for cryostat sectioning as described under “Histology and Immunostaining.” Sections were incubated in streptavidin–Alexa Fluor 750 (2 µg/mL in TBS) for 1 h at room temperature with gentle agitation, then washed three times in TBS.

##### Image analysis and modeling

Fluorescence images were analyzed using a custom MATLAB pipeline. Briefly, striatal regions of interest (ROIs) were defined within each brain section, and a multi-scale background subtraction was applied to each channel using iterative Gaussian filtering with increasing kernel sizes, producing background-subtracted intensity images. Astrocyte cell bodies were segmented using a difference-of-Gaussians approach applied to the ^NLS^dTomato channel, followed by morphological opening to remove noise. Segmented objects were filtered by size and intensity to retain only high-quality astrocyte nuclei. For each segmented cell, mean pixel intensities were extracted from the Streptavidin-750 and Alexa647^DART^ channels across both signal and background pixel populations. Per-section background-subtracted intensity values were computed by subtracting the mean background pixel intensity from the mean signal pixel intensity for each channel. To normalize each channel to its maximum, background-subtracted intensities were divided by the maximum of an independently-fit single exponential (y-intercept for Streptavidin-750, fit to y = A·exp(-k·t), plateau for Alexa647^DART^, fit to y = A·(1 – exp(-k·t))), yielding values on a 0–1 scale. Normalized fluorescence ratios were then computed as 750/(750+647) for each time point. These ratios were fit to a biphasic exponential decay model, y = A·exp(-k₁·t) + (1-A)·exp(-k₂·t), constrained such that y(0) = 1 (all surface HTP biotin-labeled), y(∞) = 0 (all surface HTP replaced by newly synthesized protein), and k₁ + k₂ = 1 h⁻¹. This constraint reduced the model to two free parameters: A (the fractional amplitude of the fast component) and k₁ (the fast rate constant), with k₂ = 1 – k₁. Model fitting was performed by nonlinear least squares (MATLAB *lsqcurvefit*). Component half-lives were calculated as t½ = ln(2)/k, and an effective half-life was defined as the time at which the fitted curve crossed y = 0.5. Uncertainty was estimated by residual bootstrap resampling (10,000 iterations), with 95% confidence intervals derived from the 2.5th and 97.5th percentiles for each fitted parameter and the effective half-life.

#### Electron Microscopy

Adult mice (8 weeks old) were stereotaxically injected with AAV.PHP.AX-GfaABC1D-HTP-NLSdTomato-4×6T (2 × 10^12^ vg/mL) into the dSTR and implanted with a guide cannula, as described above. Three weeks post-surgery, mice were head-fixed and received a unilateral intracranial infusion of 0.9 µL of 30 µM Biotin^DART^. 4 h after infusion, mice were deeply anesthetized by isoflurane inhalation in a closed chamber with physical separation from the anesthetic until cessation of breathing and loss of response to toe and tail pinch, then transcardially perfused for ultrastructural preservation. Perfusion was performed using a peristaltic pump at a slow flow rate (∼6.75 mL/min), initially with 0.1 M sodium cacodylate buffer (pH 7.4), followed by ∼30 mL freshly prepared 4% PFA in 0.075 M sodium cacodylate buffer (pH 7.4) with 2 mM CaCl_2_. Brains were dissected and post-fixed overnight at 4 °C. The following day, brains were sectioned on a vibratome, and 50 µm coronal sections containing the region of interest were collected. Free-floating sections were incubated with streptavidin-Alexa Fluor 647 FluoroNanogold (Nanoprobes) for 2 h at room temperature to visualize Biotin^DART^ labeling, washed extensively, then briefly mounted onto glass slides and coverslipped with water for rapid fluorescence imaging on a slide scanner to localize Biotin^DART^-positive regions. Coverslips were then carefully removed, and sections were transferred to a cryoprotectant solution (0.25 M sucrose, 7 mM MgCl_2_, 50% glycerol in 0.05 M sodium cacodylate buffer) and stored at−20 °C until further processing. Prior to ultrastructural processing, sections were retrieved from cryoprotectant and washed to remove glycerol. Sections selected for ultrastructural analysis were then further fixed in 2.5% glutaraldehyde in PBS, washed three times for 10 min each in 0.15 M sodium cacodylate buffer, and subsequently washed three times for 10 min each in distilled water. Regions of interest were punched out from the vibratome sections, and silver enhancement was tested at 5, 30, 45, or 60 min; 45 min provided optimal labeling and was used for all subsequent analyses. Following silver enhancement, samples were washed six times in distilled water (5 min each) and stained overnight at 4°C with 1% uranyl acetate in water. Tissues were then dehydrated through a graded ethanol series, infiltrated with resin, and polymerized. Ultrathin sections were cut perpendicular to the original vibratome sections to distinguish the positive signal from background and preserve anatomical orientation and local labeling patterns. Sections were imaged on a Thermo Fisher T12 transmission electron microscope operated at 80 kV using a Gatan OneView 4k × 4k camera. The majority of images were acquired at 13,000× magnification.

#### Ex-Vivo 2-Photon Calcium Imaging

##### Slice preparation

For 2P imaging, adult mice (8 weeks old) were stereotaxically injected with 1 μL of a mixture of AAV.PHP.AX-GfaABC1D-HTP-^NLS^dTomato-4×6T (2 × 10^12^ vg/mL) and AAV5-GfaABC1D-GCaMP6f (7×10¹² vg/mL) unilaterally into the dSTR (AP +0.5, ML −1.9, DV −2.8 and −2.3). 3−4 weeks post-injection, mice were deeply anesthetized with 200 mg/kg tribromoethanol (Avertin) and rapidly decapitated. The brain was quickly removed and placed in an aerated (95% O_2_, 5% CO_2_), ice-cold slicing solution containing: 125 mM NaCl, 2.5 mM KCl, 1.25 mM NaH_2_PO_4_, 25 mM NaHCO_3_, 10 mM glucose, 0.4 mM L-ascorbic acid, 2 mM Na-pyruvate, 3 mM MgCl_2_, and 0.1 mM CaCl_2_·2H_2_O (pH 7.3−7.4, 310 mOsm/kg). Using a razor, the cerebellum and olfactory bulbs were removed, and the cerebellum side of the brain was glued using cyanoacrylate glue (Krazy Glue, Elmer’s) onto the vibratome (VT1200S, Leica) platform. Once the glue had sufficiently dried, the brain was quickly hemisected with a razor and submerged in an ice-cold slicing solution in the vibratome chamber. Coronal dSTR slices (350 μm thick) were cut with the vibratome in the same ice-cold solution. The hemisphere containing the viral infection was collected and immediately transferred into a standard artificial cerebrospinal fluid (ACSF) solution (33 °C, continuously aerated with 95% O_2_, 5% CO_2_) containing: 125 mM NaCl, 2.5 mM KCl, 1.25 mM NaH_2_PO_4_, 25 mM NaHCO_3_, 10 mM glucose, 0.4 mM L-ascorbic acid, 2 mM Na-pyruvate, 1 mM MgCl_2_, and 1 mM CaCl_2_·2H_2_O. Slices were incubated at 33 °C for 30 min, then cooled to room temperature until imaging.

##### Image acquisition

Acute slices were transferred to a circulating perfusion bath driven by a peristaltic pump (120S, Watson-Marlow) delivering continuously aerated (95% O_2_, 5% CO_2_) ACSF maintained at 33 °C via an inline temperature controller (Perfusion Temperature Controller, Scientifica). Images were acquired on an upright two-photon microscope (Bruker) equipped with a tunable laser (InSight, Spectra Physics), with beam intensity modulated by a Pockels cell and scanning performed by linear galvanometers. Imaging was performed using an Olympus XLUMPLFLN 20× water-immersion objective (numerical aperture 1.00, working distance 2.0 mm; Olympus). Imaging was controlled by PrairieView software (Bruker). GCaMP6f was excited at 920 nm, and ^NLS^dTomato-expressing astrocytes were identified by cross-excitation of dTomato with the same 920 nm laser; emission was collected through a 1040/920 filter. Images were acquired at 512 × 512 pixels with a pixel size of 0.271 μm, dwell time of 1.2 µs, 2× optical zoom, and a frame interval of 0.606 s/frame.

For each slice, experiments proceeded as follows. For experiments measuring spontaneous activity, a 10 min baseline was recorded in standard ACSF prior to tetrodotoxin (TTX) application. TTX (0.5 µM) was added to the bath, and slices were incubated for at least 10 min to block neuronal action potentials before further recordings. For experiments not requiring a spontaneous-activity baseline, TTX (0.5 µM) was present in the bath solution before slice transfer, and slices were incubated for 10 min before imaging. Fields of view were selected in the dorsolateral striatum on the basis of ^NLS^dTomato and GCaMP6f co-expression. Following TTX equilibration, a 2 min baseline was acquired, after which AMPA (100 µM) and cyclothiazide (CTZ, 100 µM) were added to the bath; a 10 min video was acquired starting 90 s after the drug addition to account for tubing dead volume and bath equilibration. Between slices, the bath was thoroughly washed with ACSF for 10−15 min.

##### ROI-based analysis

Evoked AMPA/CTZ responses were analyzed using a manual ROI-based approach in Fiji and MATLAB. Astrocytes were manually outlined in Fiji using the polygon tool and added to the ROI manager. Mean pixel values per frame were extracted for each ROI and imported into MATLAB for further analysis. ΔF/F was calculated as (F-F_0_)/F_0_, where F_0_ was defined as the mean fluorescence in the first 30 s of the 2 min baseline acquisition, prior to drug arrival in the imaging chamber. Traces were cropped to 5 min to capture the majority of the drug response. Peak response was defined as the maximum ΔF/F value within the cropped trace. The area under the curve (AUC) was calculated by numerical integration using trapezoidal method (MATLAB function).

##### AQuA2 event detection

Spontaneous astrocytic calcium activity was analyzed using AQuA2 (*21*), an event-based detection framework that identifies spatiotemporal calcium events directly from fluorescence dynamics without predefined ROIs. Raw TIFF stacks were imported into AQuA2 (run in MATLAB R2022b) and processed using the default parameter set, which was applied uniformly across all datasets and conditions to avoid parameter-dependent bias. AQuA2’s built-in pipeline automatically performed preprocessing, spatiotemporal event segmentation, and feature quantification. Per-event features, including event duration, amplitude, frequency, rise time, and other dynamic characteristics were exported for downstream analysis in custom Python scripts.

#### Cell Type Specificity Quantification

Coronal sections from dSTR of AstroHTP-injected mice were immunostained for Sox9, NeuN, and Olig2 and imaged on an Olympus FV3000 confocal microscope using identical acquisition settings across samples. Confocal images were analyzed using ilastik-generated pixel classification maps. Briefly, an ilastik (v1.4.0) Pixel Classification project was trained on n = 5 representative training images using the default feature set. RFP⁺ nuclei were hand-annotated as the foreground class, while the background was annotated across multiple distinct images to capture variability in tissue autofluorescence and non-specific signal. The trained classifier was then applied uniformly to all images in the dataset to generate probability maps. Pixel classification maps were processed using a custom Fiji macro (Synbot (*68*)) that batch-processed all images within a user-selected directory. For each image, the probability map was thresholded into a binary mask for each class, and watershed segmentation was applied to separate adjacent or touching objects. Segmented objects were counted using the Analyze Particles function with a minimum size threshold of 70 pixels^2^ and exclusion of border-touching objects. Per-class counts and image filenames were compiled into a results table and exported as a CSV file for downstream analysis, including colocalization analyses between ^NLS^dTomato^+^ and Sox9^+^/NeuN^+^/Olig2^+^nuclei to assess cell-type specificity.

#### Open-Field Behavioral and Kinematic Analysis

##### Behavioral paradigm

Adult mice (8–16 weeks old) were stereotaxically injected with AAV.PHP.AX-GfaABC1D-HTP-NLSdTomato-4×6T (2 × 10^12^ vg/mL) into the dSTR and implanted with a guide cannula, as described above. Three weeks post-surgery, mice were head-fixed and received a unilateral intracranial infusion of 0.9 µL of either: (1) 30 µM YM90K^DART^ + 3 µM Alexa647^DART^, or (2) 30 µM YM90K^PEG36^ + 30 µM Blank^DART^ + 3µM Alexa647^DART^ (control cocktail). 1 h and 24 h after DART infusions, mice were placed in the open field for 30 min. Following the second open field session, brains were collected and processed for vibratome sections as described under “Histology and Immunostaining.”

##### Behavioral recording and pose estimation

Mice were recorded in a 12-inch-square elevated open-field arena with a transparent floor and walls, surrounded by angled mirrors. A single camera positioned beneath the arena captured five simultaneous views of the mouse (one direct view through the floor and four mirror reflections) at 90 frames per second (fps) during 30 min sessions. Three-dimensional pose estimation was performed using DANNCE (*22*) which tracked 18 body parts across all camera views. For subsequent behavioral quantification, 3D coordinates were projected onto the 2D bottom-view plane and converted from pixels to centimeters using a calibrated pixel resolution derived from known arena dimensions. A subset of tracked keypoints was used for subsequent analysis: the snout, spine midpoint (used as the body center), and tail base.

##### Tracking artifact removal

Occasional escape attempts caused mice to leave the field of view of the mirror reflections, producing transient tracking failures in which keypoints moved erratically to physically implausible positions at impossible speeds. To identify and remove these artifacts, frame-to-frame speed of the body center was computed, and frames in which speed exceeded 90 cm/s were flagged. This threshold was determined empirically from inspection of tracking data across recordings; normal mouse locomotion rarely approaches this speed. Flagged frames were expanded by a 10-frame buffer on each side (∼0.11 s at 90 fps) to account for gradual tracking degradation surrounding each event. All six coordinate channels (body center, snout, and tail base, in x and y) were set to NaN at flagged frames. This ensured that all downstream metrics, including speed, turning, trajectory-based measures, and immobility, automatically excluded artifact-contaminated data.

##### Locomotion

Instantaneous speed was calculated as the frame-to-frame Euclidean displacement of the body center (spine midpoint) multiplied by the frame rate (90 fps), yielding units of cm/s. Speed values exceeding 90 cm/s were set to NaN as an additional safeguard against residual tracking noise propagated from frames adjacent to those excluded during artifact removal. Total distance traveled was computed within each 5 min bin by summing instantaneous displacements, excluding NaN frames; six bins were generated per 30 min session.

##### Turning analysis

The instantaneous heading direction of the mouse was defined as the angle from the tail base to the snout in the 2D plane. This heading angle was unwrapped to remove discontinuities at the ±180° boundary and low-pass filtered to reduce high-frequency noise. A complete turn was defined as a cumulative heading change of ≥360° in a single direction without returning through the starting angle. To guard against spurious detections, candidate turns were rejected if the heading reversed by more than 180° from the cumulative turn trajectory before completing the 360° rotation, indicating the mouse changed direction mid-turn, or if any frame between the start and completion of the candidate turn contained a NaN (i.e., a tracking artifact). Turns were classified as leftward (counterclockwise from the perspective of the mouse) or rightward (clockwise from the perspective of the mouse) based on the sign of the cumulative heading change. Net turns per session were computed as the difference between rightward and leftward turn counts; total leftward and rightward turn counts were tallied separately.

##### Thigmotaxis

To quantify thigmotaxis, the proportion of time spent in the center versus the edge of the arena was computed from the body center trajectory. The center zone was a 12.5 × 12.5 cm square region centered on the midpoint between the 1^st^ and 99^th^ percentiles of the body center’s x and y coordinates. This data-driven centering approach accounted for minor variations in tracking coordinate offsets across sessions; deviations greater than 10% of arena width from the geometric arena center were overridden by the geometric center. Center occupancy was expressed as the percentage of total frames in which the body center fell within the center zone; edge occupancy was the complement. Center occupancy was computed per 30 min session.

##### Immobility

Immobility was assessed using a bout-based analysis of smoothed locomotor speed. The instantaneous speed signal was smoothed with a 90-frame (1 s) moving median filter and log10-transformed. Frames in which the transformed speed fell below a threshold of 0 (corresponding to a raw speed of <1 cm/s) were classified as immobile. Contiguous immobile frames were grouped into bouts, and bouts shorter than 1 s (90 frames) were discarded to exclude brief pauses during active locomotion. From the remaining bouts, the total percent time immobile was computed per 30 min session.

#### Electrophysiology: Astrocyte-Targeted DART

##### Slice preparation

Young adult mice (6−8 weeks old) of either Drd1a-Cre or Adora2a-Cre lines were stereotaxically injected with 1 μL of a mixture of AAV.PHP.AX-GfaABC1D-HTP-^NLS^dTomato-4×6T (2 × 10^12^ vg/mL) and AAV5-hSyn-DIO-GFP (2.3 × 10^12^ vg/mL) unilaterally into the dSTR (AP +0.5, ML −1.9, DV −2.8 and −2.3). 3-4 weeks post-injection, mice were deeply anesthetized with 200 mg/kg tribromoethanol (Avertin) and rapidly decapitated. Brains were quickly removed and placed in ice-cold, carbogenated (95% O2, 5% CO2) sucrose-based cutting solution containing (in mM): 3.3 KCl, 1.3 NaH_2_PO_4_, 26 NaHCO_3_, 11 D-glucose, 211 sucrose, 0.5 CaCl_2_, and 10 MgCl_2_. Coronal slices (300 µm) containing the dSTR were cut on a vibratome (VT1000S, Leica). Slices were transferred to carbogenated ACSF and allowed to recover at 33 °C for 50 min, then for an additional 10 min at room temperature before recording. ACSF contained (in mM): 124 NaCl, 3.3 KCl, 1.3 NaH_2_PO_4_, 26 NaHCO_3_, 11 D-glucose, 2 CaCl_2_, and 1 MgCl_2_, and was continuously equilibrated with 95% O_2_ / 5% CO_2_ during recovery and recording. Slices were used for up to 4 h after the end of the recovery period.

##### DART Application

To inhibit astrocytic AMPARs in acute slices, YM90K^DART^ was bath-applied during recording. Slices were continuously perfused with ACSF containing either 300 nM YM90K^DART^ or a control cocktail of 300 nM YM90K^PEG36^ (an untetherable variant lacking the HaloTag ligand) and 300 nM Blank^DART^ (a pharmacologically inert tethered ligand). In both conditions, 30 nM Alexa647^DART^ was included as a fluorescent tracer to verify ligand capture. For experiments measuring miniature EPSCs (mEPSC), slices were preincubated in the corresponding DART solution for at least 30 min before recording. For experiments measuring evoked EPSCs (eEPSC), a stable 5 min baseline was first acquired in standard ACSF, after which the DART solution was bath-applied for 15 min, followed by a 10 min washout period during which recordings continued.

##### Whole-cell recordings

Whole-cell voltage-clamp recordings were obtained from GFP⁺ SPNs adjacent to ^NLS^dTomato⁺ astrocytes in a submerged recording chamber superfused with warmed ACSF (29–32 °C). Patch pipettes (2–4 MΩ) were pulled from borosilicate glass. For mEPSC recordings, pipettes were filled with an internal solution containing (in mM): 145 CsCl, 5 NaCl, 10 HEPES, 10 EGTA, 4 Mg-ATP, and 0.3 Na-GTP. For eEPSC recordings, the internal solution contained (in mM): 130 CsMeSO_4_, 5 TEA-Cl, 8 NaCl, 10 HEPES, 0.5 EGTA, 4 Mg-ATP, 0.4 Na-GTP, 10 phosphocreatine disodium salt hydrate, and 3 QX-314. All internal solutions were adjusted to pH of 7.3–7.4 with CsOH and to an osmolality of 280–285 mOsm/kg. After formation of a giga-ohm seal and break-in, cells were voltage-clamped at −70 mV. Electrical stimulation (0.1 ms pulses) for evoked recordings was delivered every 20 s through a concentric bipolar electrode (FHC). For each cell, stimulus intensity was adjusted to evoke approximately 40% of the maximal eEPSC amplitude by gradual increase from threshold. Paired-pulse ratios (PPRs) were calculated as the amplitude of the second EPSC divided by that of the first EPSC, using a 50 ms interstimulus interval. Access resistance and holding current were monitored throughout each recording, and cells exhibiting instability were excluded from analysis.

For single pulse eEPSC recordings, stimuli were delivered every 20s. For paired-pulse recordings, stimulus pairs were delivered every 30s, with a 50-ms interstimulus interval (ISI) within each pair.

AMPAR component was measured as the peak current at −70 mV, whereas the NMDAR component was measured at +40 mV at the post-stimulus latency at which the AMPAR-mediated current recorded at −70 mV had decayed to baseline.

##### Pharmacology

For mEPSC recordings, ACSF contained tetrodotoxin (TTX, 1µM) to block action potentials and gabazine (SR-95531, 10 µM) to block GABA_A_ receptor-mediated currents. For eEPSC recordings, ACSF contained gabazine to isolate excitatory transmission. To isolate AMPAR-mediated currents in experiments testing astrocyte selectivity, the NMDAR antagonist CPP (10 µM) was included in the bath. In experiments testing the requirement for ionotropic NMDAR conductance, the open-channel NMDAR blocker MK-801 (10 µM) was applied to the bath. To assess the contribution of NMDAR co-agonist availability, D-serine (10 µM) was co-applied with MK-801. To enzymatically deplete extracellular D-serine, D-amino acid oxidase (DAAO, 0.3 U/mL) was bath-applied together with catalase (300 U/mL) using the same recording paradigm as for eEPSCs. All drugs were applied via the perfusion line and allowed to equilibrate in the recording chamber prior to or during recording, as described for each experiment in the results.

##### Data acquisition and analysis

Signals were acquired using a HEKA EPC 10 amplifier controlled by PATCHMASTER NEXT software (HEKA Elektronik). mEPSCs were recorded for 5 min per cell following a brief stabilization period after break-in. eEPSCs were recorded for up to 30 min per cell, and slices used for eEPSC recordings were not reused. For mEPSC quantification, the most stable 100 s segment from each recording was selected for analysis. Event detection and measurements were performed in FITMASTER NEXT (HEKA Elektronik) using consistent detection thresholds and criteria across cells. For eEPSC quantification, peak-to-anti-peak amplitudes were measured in PATCHMASTER NEXT. Representative traces were exported from FITMASTER NEXT and prepared for figures in Inkscape.

#### Electrophysiology: Neuron-Targeted DART

##### Slice preparation

Adora2a-Cre mice (8–10 weeks old) were stereotaxically injected with AAV10-CAG-DIO-HTP-2A-dTomato-WPRE (active HTP, 1.5 × 10^12^ vg/mL) or AAV10-CAG-DIO-ddHTP-2A-dTomato-WPRE (dead HTP control, 1.5 × 10^12^ vg/mL) at 80 nL per depth across multiple sites in the dSTR (AP +0.05 mm, ML ±1.7 mm, DV −2.8, −2.3, and −1.8 mm; AP +0.15 mm, ML ±2.5 mm, DV −3.1, −2.76, and −2.1 mm; AP +0.85 mm, ML ±1.3 mm, DV −3.0, −2.5, and −2.0 mm; and AP +0.95 mm, ML ±2.1 mm, DV −3.1, −2.6, and −2.1 mm, all relative to bregma). After 3–5 weeks for expression, mice were deeply anesthetized with isoflurane and decapitated. Coronal brain slices (300 µm) containing dSTR were cut on a Vibratome (Leica, VT1200S), in ice-cold high sucrose cutting solution containing (in mM): 220 sucrose, 3 KCl, 1.25 NaH2PO4, 25 NaHCO_3_, 12 MgSO_4_.7H_2_O, 10 glucose, and 0.2 CaCl_2_, bubbled with 95% O_2_ / 5% CO_2_. Slices were transferred to ACSF and allowed to recover at 33 °C for 40–60 min, then cooled to room temperature (22–24 °C) before recording. ACSF contained (in mM): 120 NaCl, 3.3 KCl, 1.23 NaH_2_PO_4_, 1 MgSO_4_, 2 CaCl_2_, 25 NaHCO_3_, and 10 glucose (pH 7.3), continuously equilibrated with 95% O_2_ / 5% CO_2_.

##### DART Application

To test whether YM90K^DART^ inhibits neuronal AMPARs when tethered to neuronal HTP, the ligand was bath-applied to slices expressing either active HTP or catalytically inactive HTP (ddHTP, negative control). Slices were continuously perfused with ACSF containing 300 nM YM90K^DART^; 30 nM Alexa647^DART^ was included as a fluorescent tracer to verify ligand capture. A stable 7.5 min baseline was first acquired in standard ACSF, after which the DART solution was bath-applied for 15 min, followed by a 7.5 min washout period during which recordings continued.

##### Whole-cell recordings

Whole-cell voltage-clamp recordings were obtained from dTomato^+^ SPNs in slices perfused with bubbled ACSF at 29–30 °C (2 mL/min). Patch pipettes (4–6 MΩ) were filled with an internal solution containing (in mM) 130 CsMeS, 1 MgCl_2_, 0.5 EGTA, 10 HEPES, 4 Mg-ATP, 0.5 Na-GTP, 10 Na_2_-phosphocreatine, and 4 QX-314, adjusted to pH 7.3 with CsOH and 290 mOsm. Cells were voltage clamped at – 70 mV. Electrical stimulation (0.3 ms pulses, 60–70% maximum response, 150–300 µA) was delivered every 30s through a stimulating electrode placed 60–100 µm from the recorded neuron. Series resistance was monitored using 5–10 mV hyperpolarizing steps interleaved with stimuli, and cells with series resistance changes greater than ∼15% during the experiment were excluded.

##### Pharmacology

To isolate AMPAR-mediated EPSCs, picrotoxin (50 µM, GABA_A_ receptor antagonist) and APV (10 µM, NMDAR antagonist) were included in the bath.

##### Data acquisition and analysis

Signals were acquired using a Multiclamp 700B amplifier (Molecular Devices), filtered at 10 kHz, and digitized using an Axon Digidata 1550B with pClamp 10.7 (Molecular Devices). Data were analyzed in Clampfit 10.7.

#### Quantification and statistical analysis

Reported values are mean ± SEM. Statistical analyses were performed in GraphPad Prism 10. Comparisons used Student’s t-test (paired or unpaired), the Mann–Whitney test, the Wilcoxon matched-pairs signed-rank test, two-way ANOVA (with Fisher’s LSD or Šídák’s multiple-comparisons post hoc tests, as indicated), and the Kruskal–Wallis test (with Dunn’s post hoc test), as specified in each figure legend. The statistical test, sample size, and P value are reported in each figure or figure legend. No statistical methods were used to predetermine sample size.

For *ex vivo* two-photon calcium imaging, multiple regions of interest (ROIs) or calcium events were recorded per slice and per animal; to avoid pseudoreplication, these measurements were aggregated at the slice level, and statistical comparisons were performed at the slice level as the unit of analysis. ROI, event, slice, and animal numbers are reported in the corresponding figure legends.

For two-photon experiments, conditions being compared (for example, Blank^DART^ versus YM90K^DART^) were interleaved and acquired side by side, using cage mates where possible, with groups balanced to minimize confounding differences. In vivo experiments used male mice only; where possible, cage mates were used, and groups were balanced to minimize confounding factors.

For whole-cell electrophysiology, the recorded cell was treated as the statistical unit, following standard practice for slice electrophysiology; cells were sampled across multiple mice for each condition, and the numbers of cells and mice are reported in each figure legend.

#### Antibodies

| Reagent or Resource | Source | Identifier |
| --- | --- | --- |
| Rabbit polyclonal anti-Sox9 | Millipore | Cat# AB5535;<br>RRID:AB_2239761 |
| Mouse Monoclonal anti-NeuN<br>Antibody, clone A60 | Millipore | Cat# MAB377,<br>RRID:AB_2298772 |
| Mouse IgG2a monoclonal anti-Olig2,<br>clone 211F1.1 | Millipore | Cat# MABN50,<br>RRID:AB_10807410 |
| Streptavidin–FluoroNanogold 647 | FluoroNanogold /<br>Nanoprobes | Cat# 7516 |
| Goat anti-mouse IgG2a Alexa Fluor<br>568 | Thermo Fisher Scientific | RRID:AB_2535773 |
| Goat anti-mouse IgG2b Alexa Fluor<br>568 | Thermo Fisher Scientific | RRID:AB_2535780 |
| Goat anti-mouse Alexa Fluor 488 | Thermo Fisher Scientific | RRID:AB_2534057 |
| Goat anti-rabbit Alexa Fluor 568 | Thermo Fisher Scientific | RRID:AB_143157 |
| Goat anti-rabbit Alexa Fluor 647 | Thermo Fisher Scientific | RRID:AB_2921064 |
| Donkey anti-mouse Alexa Fluor 647 | Jackson<br>ImmunoResearch | RRID:AB_162542 |
| Donkey anti-rabbit Alexa Fluor 488 | Jackson<br>ImmunoResearch | RRID:AB_3107111 |

#### Chemicals, Peptides, and Drugs

| Reagent or Resource | Source | Identifier |
| --- | --- | --- |
| YM90K<sup>PEG36</sup> | Tadross Lab | Custom synthesis |
| YM90K<sup>DART</sup> | Tadross Lab | Custom synthesis |
| Blank<sup>DART</sup> | Tadross Lab | N/A |
| Alexa647<sup>DART</sup> | Tadross Lab | N/A |
| Biotin<sup>DART</sup> | Tadross Lab | N/A |
| D-Serine | Sigma-Aldrich | Cat# S4250-5 |
| RS-AMPA | Tocris Bioscience | Cat# 0169 |
| Cyclothiazide | Tocris Bioscience | Cat# 0713 |
| NBQX | Tocris Bioscience | Cat# 0373 |
| D-APV (AP5) | Tocris Bioscience | Cat# 0106/1 |
| (R)-CPP | Tocris Bioscience | Cat# 0247 |
| (+)-MK 801 maleate | Tocris Bioscience | Cat# 0924 |
| Picrotoxin | Tocris Bioscience | Cat# 1128/1G |
| Tetrodotoxin (TTX) | Tocris Bioscience | Cat# 1078/1 |
| Gabazine (SR-95531) | Tocris Bioscience | Cat# 1262 |
|  |  | Cat# A5222-100UN |
| D-Amino Acid Oxidase (DAAO; porcine kidney) | Sigma-Aldrich |  |
| Catalase from bovine liver | Sigma-Aldrich | Cat# C1345-1G |

#### Experimental Models: Organisms/Strains

| Reagent or Resource | Source | Identifier |
| --- | --- | --- |
| C57BL/6J mice | Jackson Laboratory | RRID:IMSR_JAX:000664 |
| Drd1a-Cre mice | GENSAT / Jackson Laboratory | RRID:IMSR_JAX:028298 |
| Adora2a-Cre mice | MMRRC / Jackson Laboratory | RRID:IMSR_JAX:040186 |

#### Cell Lines

| Reagent or Resource | Source | Identifier |
| --- | --- | --- |
| One Shot™ Stbl3™ Chemically Competent E. coli | Thermo Fisher Scientific | Cat# C737303 |
| HEK293T cells | ATCC | RRID:CVCL_0063 |

#### Recombinant DNA / Viral Vectors

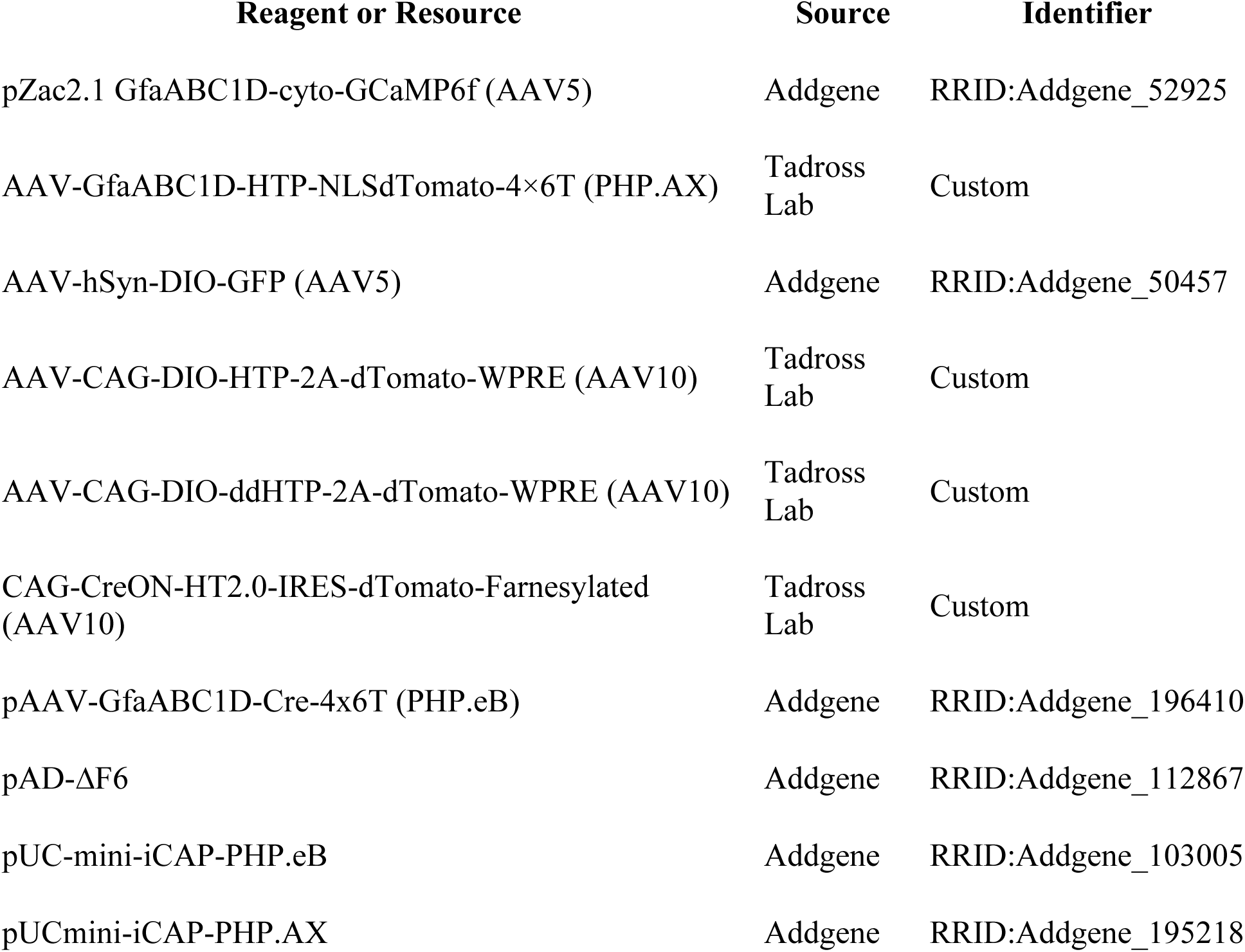

| Reagent or Resource | Source | Identifier |
| --- | --- | --- |
| pZac2.1 GfaABC1D-cyto-GCaMP6f (AAV5) | Addgene | RRID:Addgene_52925 |
| AAV-GfaABC1D-HTP-NLSdTomato-4×6T (PHP.AX) | Tadross Lab | Custom |
| AAV-hSyn-DIO-GFP (AAV5) | Addgene | RRID:Addgene_50457 |
| AAV-CAG-DIO-HTP-2A-dTomato-WPRE (AAV10) | Tadross Lab | Custom |
| AAV-CAG-DIO-ddHTP-2A-dTomato-WPRE (AAV10) | Tadross Lab | Custom |
| CAG-CreON-HT2.0-IRES-dTomato-Farnesylated (AAV10) | Tadross Lab | Custom |
| pAAV-GfaABC1D-Cre-4x6T (PHP.eB) | Addgene | RRID:Addgene_196410 |
| pAD-ΔF6 | Addgene | RRID:Addgene_112867 |
| pUC-mini-iCAP-PHP.eB | Addgene | RRID:Addgene_103005 |
| pUCmini-iCAP-PHP.AX | Addgene | RRID:Addgene_195218 |

#### Software and Algorithms

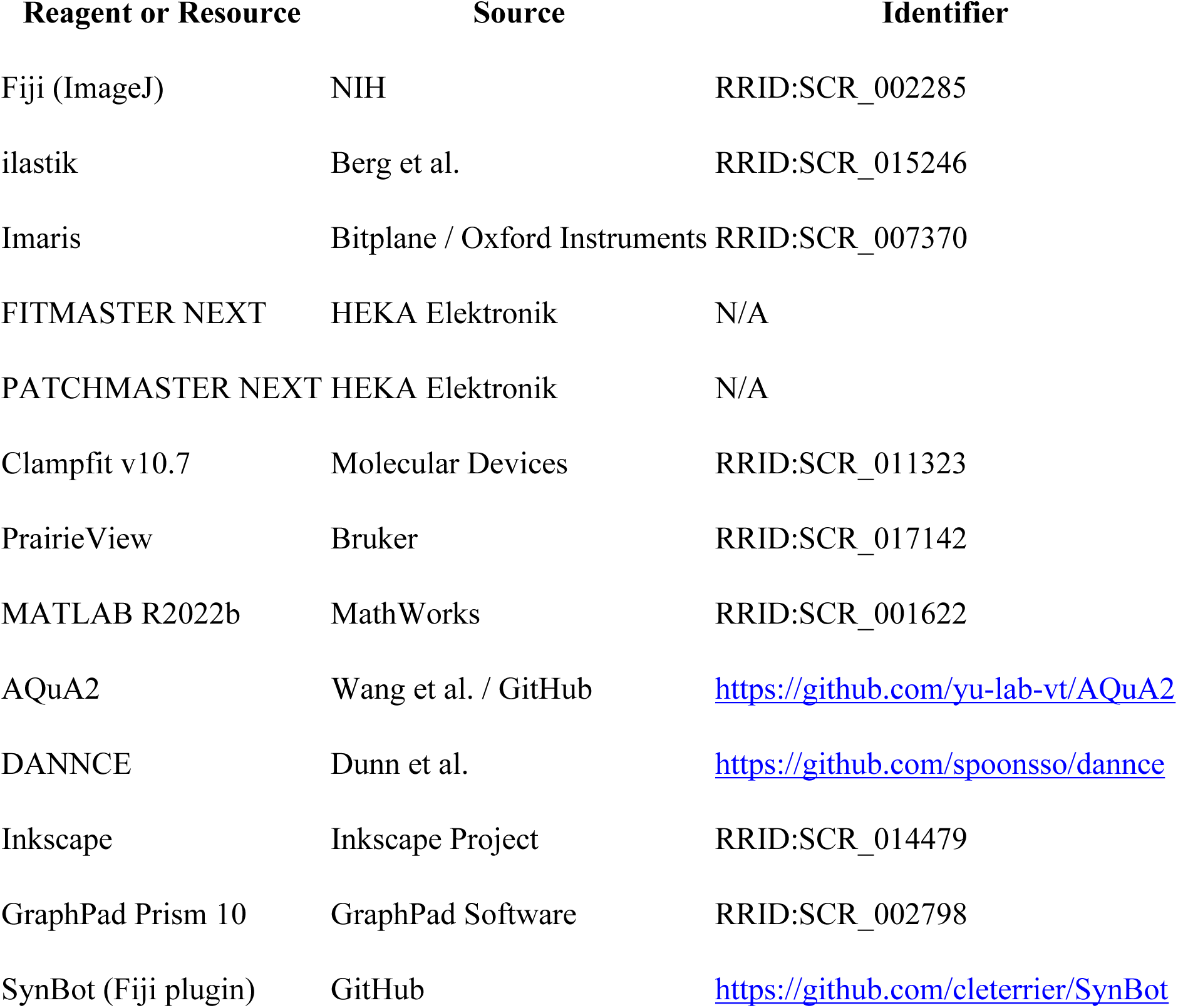

#### Other

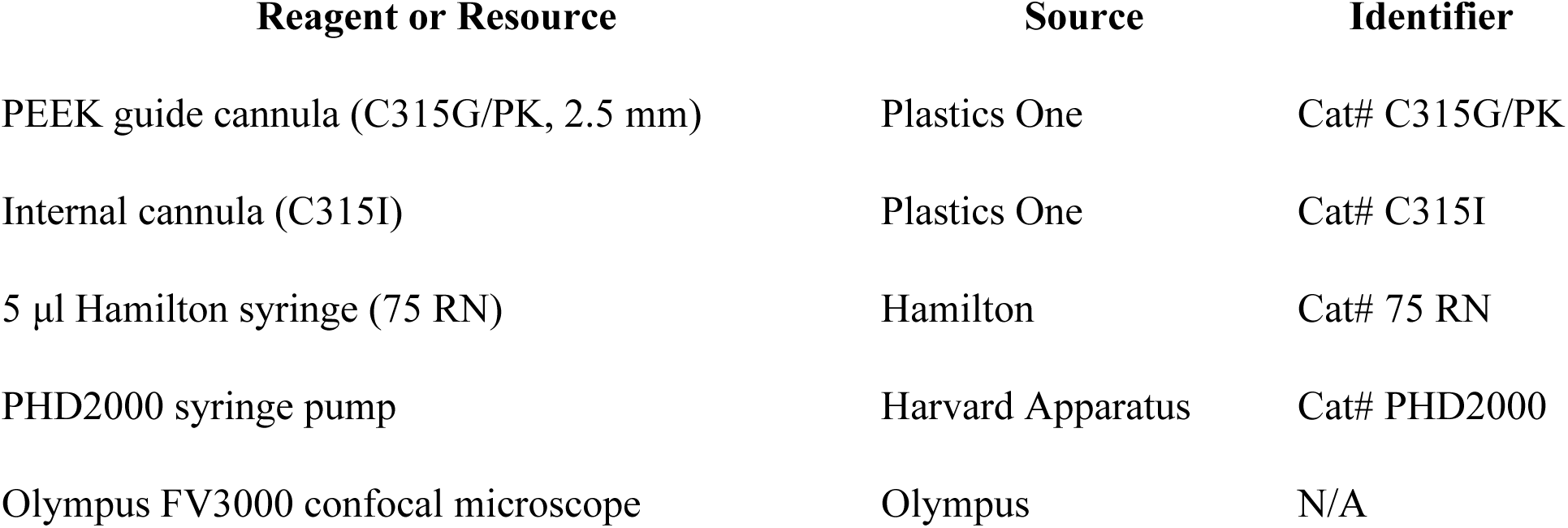
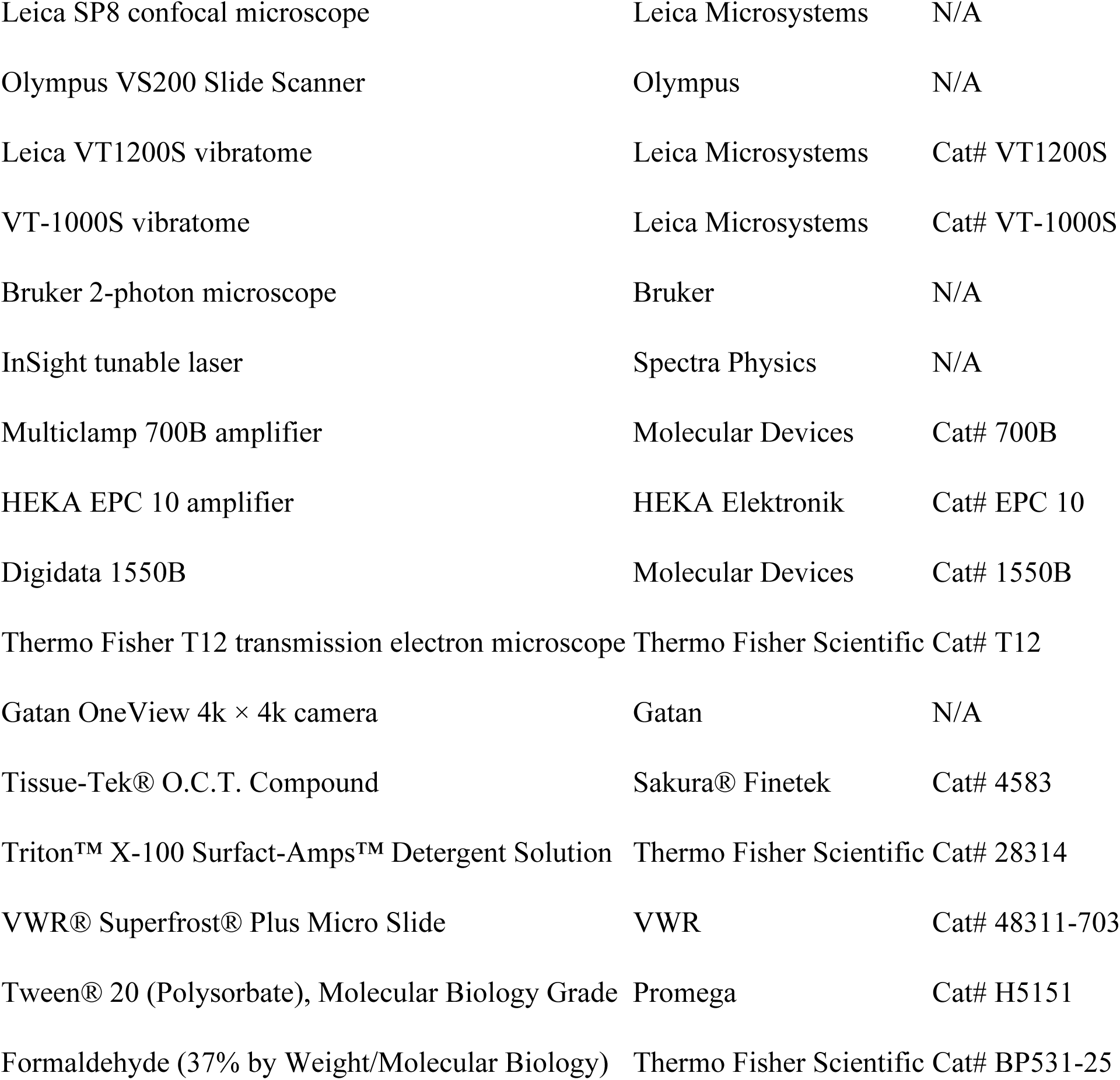

**Figure S1. Supplement to Figure 1**. **Further characterization of astrocyte-specific DART. (A)** AstroHTP-transduced striatal astrocytes capture DART ligands at perisynaptic processes. AstroHTP was injected into the dSTR of adult C57BL/6J mice. Three weeks post-injection, 30 µM biotin-conjugated HTP ligand (Biotin^DART^) was infused into the striatum of head-fixed mice for 15 min. Coronal striatal sections were processed for immuno-electron microscopy and labeled with Alexa Fluor 647-conjugated FluoroNanogold™ to visualize HTP-bound DART ligands. Representative image from a coronal section of a C57BL/6J mouse brain expressing AstroHTP (magenta), 4 h post-infusion with Biotin^DART^ and post-hoc labeling with Alexa Fluor 647-conjugated FluoroNanogold™ (grey). Magenta box, negative-control hemisphere (no virus injected); blue box, experimental hemisphere. Scale bars: 1000 µm, 10 µm.(**B)** Representative electron micrograph from a coronal striatal section 4 h after intracranial infusion of Biotin^DART^, showing Alexa647-nanogold labeling of astrocytic processes (green). Blue outline, experimental hemisphere; magenta outline, control hemisphere. Scale bar, 100 nm. **(C–E)** AstroHTP can be used to target astrocytes in different brain regions. **(C)** AstroHTP was injected into the motor cortex or the hippocampus of adult C57BL/6J mice. Three weeks post-injection, 3 µM Alexa647-DART was infused into the ventricles of head-fixed mice for 15 min. **(D)** Representative images from a coronal section of an adult C57BL/6J mouse hippocampus and motor cortex, 24 h post-infusion with Alexa647^DART^, showing ligand capture. DAPI (blue), dTomato-NLS (magenta), Alexa647^DART^ (grey). Scale bar, 1000 µm. **(E)** Specificity of AstroHTP in the hippocampus: coronal sections were immunolabeled with antibodies against cell-type-specific markers: SOX9 (astrocytes), NeuN (neurons), and OLIG2 (oligodendrocytes). Scale bar, 100 µm.

**Figure S2. Supplement to Figure 2**. **Further characterization of astrocyte-specific AMPAR attenuation with YM90K^DART^. (A–B)** Astrocytes express AMPAR subunits. Summary of published TRAP RNA-seq (*17*) **(A)** and proteomic (*18*) **(B)** datasets reporting glutamate receptor expression in striatal astrocytes. **(C)** YM90K^DART^-mediated attenuation of AMPAR function at low ambient drug concentrations. The DART tether (blue line) anchors the pharmacological ligand to HTP on target cells, selectively enriching the drug at the cell surface and enabling effective receptor inhibition despite minimal free ligand in the surrounding tissue. Tethered pharmacology is governed by two separable parameters: CC50 (cumulative capture 50%), which defines the exposure (concentration × time, nM·min) required for half-maximal tethered-drug effect on +HTP cells and reflects irreversible HaloTag ligand-driven capture; and AC50 (ambient concentration 50%), which defines the ambient concentration (nM) producing half-maximal reversible drug effect on −HTP cells and is determined by the pharmacological moiety. Consequently, non-HTP-expressing cells (grey outline) experience only low ambient drug levels and remain largely unaffected. Adapted from Shields et al (*11*). **(D–F)** Astrocyte Ca²⁺ response to AMPA application. **(D)** *Upper:* two-photon *ex vivo* Ca²⁺ imaging paradigm in astrocytes. Dorsal striatal (dSTR) astrocytes from adult C57BL/6J mice were co-transduced with AstroHTP and GCaMP6f. Cyclothiazide (CTZ) was applied to block AMPAR desensitization and enhance astrocytic Ca²⁺ responses, and tetrodotoxin (TTX) was applied to block action potential-dependent neuronal activity, isolating the direct astrocytic response to AMPAR activation. *Lower:* representative images of GCaMP6f-expressing astrocytes with ROIs selected for analysis, and corresponding ΔF/F raster plots showing Ca²⁺ activity of each ROI throughout the experiment. **(E)** ΔF/F traces showing astrocytic Ca²⁺ responses to AMPA (100 µM) in the presence of CTZ (100 µM) and TTX (0.5 µM), with or without NBQX. Traces correspond to ACSF control (grey) or NBQX (orange). Shading indicates mean ± SEM. **(F)** NBQX reduced peak Ca²⁺ responses (ACSF 1.08 ± 0.196; NBQX 0.0291 ± 0.00609; unpaired t-test, P = 0.0063; N=3 mice, n = 4 slices (ACSF) and 3 slices (NBQX); 34 and 25 ROIs, respectively). **(G)** Astrocytic responses to AMPA are attenuated by YM90K^DART^. *left:* two-photon *ex vivo* Ca²⁺ imaging paradigm in astrocytes. Dorsal striatal (dSTR) astrocytes from adult C57BL/6J mice were co-transduced with AstroHTP and GCaMP6f. Slices were pre-incubated with DART ligands before application of AMPA with cyclothiazide (CTZ). Tetrodotoxin (TTX) was applied to block action potential-dependent neuronal activity, isolating the direct astrocytic response to AMPAR activation. *right:* representative images of GCaMP6f-expressing astrocytes with ROIs selected for analysis, and corresponding ΔF/F raster plots showing Ca²⁺ activity of each ROI throughout the experiment.

**Figure S3 Supplementary for Figure 3**. **Turning Behavior and Additional Locomotor Parameters 24 Hours Post DART-Infusion (A)** Cumulative net turns of the mouse, showing both right and left turns over time and their accumulation over the session. Data are mean ± SEM. N = 10 control, 11 YM90K^DART^-infused mice. **(B)** Fraction of right turns. Control 0.801± 0.0743; YM90K^DART^ 0.628± 0.0718; Mann– Whitney test, P = 0.1517; N = 10 control, 11 YM90K^DART^-infused mice. **(C)** Turn-diameter histogram (1 cm bins) 24 h after control (grey) or YM90K^DART^ infusion. Left turns are plotted as downward lines and right turns as upward lines. Data are mean ± SEM. N = 10 control, 11 YM90K^DART^-infused mice. **(D)** Total turns at 24 h post-DART infusion. Mann-Whitney test, P = 0.3494. **(E)** Both groups showed a reduction in distance traveled between 1 h and 24 h post-infusion (two-way ANOVA, main effect of timepoint: F(1,19) = 14.99, P = 0.0010), with no effect of treatment (two-way ANOVA, main effect of treatment: F(1,19) = 0.05, P = 0.8237). **(F)** % Time in center did not differ between control and YM90K^DART^-treated mice or between time points (two-way ANOVA, main effect of treatment: F(1,19) = 0.6397, P = 0.4337, or effect of time F (1, 19) = 0.1260, P = 0.726. **(G)** Time spent immobile did not differ between groups (two-way ANOVA, main effect of treatment: F(1,19) = 0.08, P = 0.7825) but decreased between 1 h and 24 h in both groups (two-way ANOVA, main effect of timepoint: F(1,19) = 10.72, P = 0.0040).

**Figure S4 Supplementary for Figure 3**. **Neuronal AMPAR inhibition of iSPNs via YM90K^DART^ is not NMDAR dependent. (A**) Adora2a-Cre mice (iSPNs) were transduced with AAV-CAG-DIO-HTP-2A-dTomato-WPRE or a binding-deficient control, AAV-CAG-DIO-^dd^HTP-2A-dTomato-WPRE, which cannot capture the HaloTag ligand and therefore leaves YM90K^DART^ untethered. AMPA-R mediated eEPSC were recorded from striatal slices in the ACSF solution containing picrotoxin (50 µM) and APV (10 µM), and bath-applied with 300 nM YM90K^DART^/30 nM Alexa647^DART^ after a 7.5-minute baseline. Recordings were obtained from neurons expressing HTP or ^dd^HTP (control). **(B)** Representative image of iSPNs expressing HTP (red), capturing Alexa647DART (grey). Scale bar, 10 µm. **(C)** Time course of normalized eEPSC amplitude showing a marked reduction following YM90K^DART^ treatment in HTP-expressing cells (red) compared with ^dd^HTP-expressing cells (grey). Data are mean ± SEM. ^dd^HTP condition: 8 cells from 4 mice, ^+^HTP condition: 7 cells from 3 mice. **(D)** Representative eEPSC traces at baseline (black) and after YM90K^DART^ application (red), shown for both ^dd^HTP and ^+^HTP cells. **(E)** eEPSC amplitude was significantly reduced following YM90K^DART^ treatment (control 87.4 ± 4.45%; YM90K^DART^ 30.2 ± 8.37%; Mann– Whitney test, P = 0.0003; 17.5–22.5 min). ^dd^HTP condition: 8 cells from 4 mice, ^+^HTP condition: 7 cells from 3 mice.

**Figure S5 Supplementary for Figure 3**. **Astrocytic AMPAR inhibition of SPNs does not alter basal synaptic transmission. (A)** Experimental design for mEPSC recordings. **(B)** Representative recording configuration showing a GFP-positive dSPN (green) patched adjacent to an NLS-dTomato-positive astrocyte (red). Scale bar, 10 µm. **(C)** Distance between patched dSPNs and neighboring astrocytes did not differ between groups (control 9.67 ± 1.25 µm; YM90K^DART^ 8.17 ± 0.852 µm; Mann–Whitney test, P = 0.4463; control n = 15 cells from 4 mice; YM90K^DART^ n = 16 cells from 4 mice. **(D)** Representative mEPSC traces recorded from dSPNs in control (grey) and YM90K^DART^ treated (blue) conditions. **(E)** mEPSC amplitude in dSPNs was unchanged (control 10.2 ± 0.345 pA; YM90K^DART^ 9.70 ± 0.288 pA; Mann–Whitney test, P = 0.4008; control n = 15 cells from 4 mice; YM90K^DART^ n = 16 cells from 4 mice). **(F)** mEPSC frequency in dSPNs was unchanged (control 0.897 ± 0.129 Hz; YM90K^DART^ 1.31 ± 0.296 Hz; Mann–Whitney test, P = 0.8609; control n = 15 cells from 4 mice; YM90K^DART^ n = 16 cells from 4 mice). **(G)** Representative recording configuration showing a GFP-positive iSPN (green) patched adjacent to an NLS-dTomato-positive astrocyte (red). Scale bar, 10 µm **(H)** Distance between patched iSPNs and neighboring astrocytes did not differ between groups (control 8.29 ± 0.987 µm; YM90K^DART^ 8.23 ± 1.07 µm; Mann– Whitney test, P = 0.9668; control n = 20 cells from 4 mice; YM90K^DART^ n = 19 cells from 4 mice). **(I)** Representative mEPSC traces recorded from iSPNs in control (grey) and YM90K^DART^ treated (blue) conditions. **(J)** mEPSC amplitude in iSPNs was unchanged (control 9.12 ± 0.258 pA; YM90K^DART^ 9.65 ± 0.244 pA; Mann–Whitney test, P = 0.1841; control n = 20 cells from 4 mice; YM90K^DART^ n = 19 cells from 4 mice). **(K)** mEPSC frequency in iSPNs was unchanged (control 0.399 ± 0.0413 Hz; YM90K^DART^ 0.427 ± 0.0435 Hz; Mann–Whitney test, P = 0.3803; control n = 20 cells from 4 mice; YM90K^DART^ n = 19 cells from 4 mice).

**Figure S6 Supplementary for Figures 4 and 5**. **Astrocyte AMPAR Blockade Reduces eEPSCs via NMDAR-Dependent, D-Serine-Sensitive Mechanism. (A)** Comparison of the change in eEPSC amplitude after 10 min of stimulation (ΔeEPSC, % of baseline): continuous −21.7 ± 8.62%; paused −37.2 ± 13.1% (Mann–Whitney test, P = 0.4234; continuous n = 9 cells from 5 mice, paused n = 8 cells from 4 mice). **(B)** Time course of the first eEPSC amplitude in the PPR experiment, normalized to baseline. YM90K^DART^ treatment reduced eEPSC amplitude.). Both measures: n = 9 cells from 5 mice. **(C)** Time course and quantification of PPR, showing no change following YM90K^DART^ treatment. **(D)** Paired comparison of eEPSC amplitude at baseline and after 10 min of YM90K^DART^ application, 64.4% ± 8.16 % (Wilcoxon test, P = 0.0034), n=12 cells, 10 mice. **(E)** Paired comparison of eEPSC amplitude at baseline and 10–20 min post-infusion of YM90K^DART^ and MK-801, 80.3%± 7.62% (Wilcoxon test, P = 0.0134), n=13 cells, 9 mice. **(F)** Paired comparison of eEPSC amplitude at baseline and 10–20 min post-infusion of YM90K^DART^ and MK-801+D-serine, 108%± 5.58% (Wilcoxon test, P = 0.2061), n=11 cells, 9 mice. **(G)** Paired comparison of eEPSC amplitude at baseline and 10–20 min post-infusion of vehicle, 105%± 9.96% (Wilcoxon test, P = 0.6377), n=11cells, 7 mice. **(H)** Paired comparison of eEPSC amplitude at baseline and 10–20 min post-infusion of DAAO, 72.7%± 7.40% (Wilcoxon test, P = 0.0040), n=14 cells, 8 mice.

**Supplementary Videos: 2P imaging of astrocyte calcium transients in response to AMPA bath application.**

Supp Video 1. Representative time-lapse video showing astrocyte response to application of 100uM AMPA+100uM CTZ

Supp Video 2. Representative time-lapse video showing astrocyte lack of response to application of 100uM AMPA+100uM CTZ pre-incubated in 30uM NBQX

Supp Video 3. Representative time-lapse video showing astrocyte response to application of 100uM A MPA+100uM CTZ pre-incubated in 1uM Blank ^DART^.

Supp Video 4. Representative time-lapse video showing astrocyte lack of response to application of 100uM A MPA+100uM CTZ pre-incubated in 1uM YM90K^DART^.

**Supplementary Videos: 2P imaging of astrocyte spoteneuos calcium transients**

Supp Video 5. Representative time-lapse video showing AquA2 analysis of spoteneuos calciuym events in slices pre-incubated in 1uM Blank ^DART^.

Supp Video 6. Representative time-lapse video showing AquA2 analysis of spoteneuos calcium events in slices pre-incubated in 1uM YM90K^DART^.

Supp Video 7. Representative time-lapse video showing DANNCE tracking of mice in open field.

