## Supplementary figures and images for "An astrocytic AMPAR-dependent parity signal preserves synaptic strength"

### Supplementary Figures

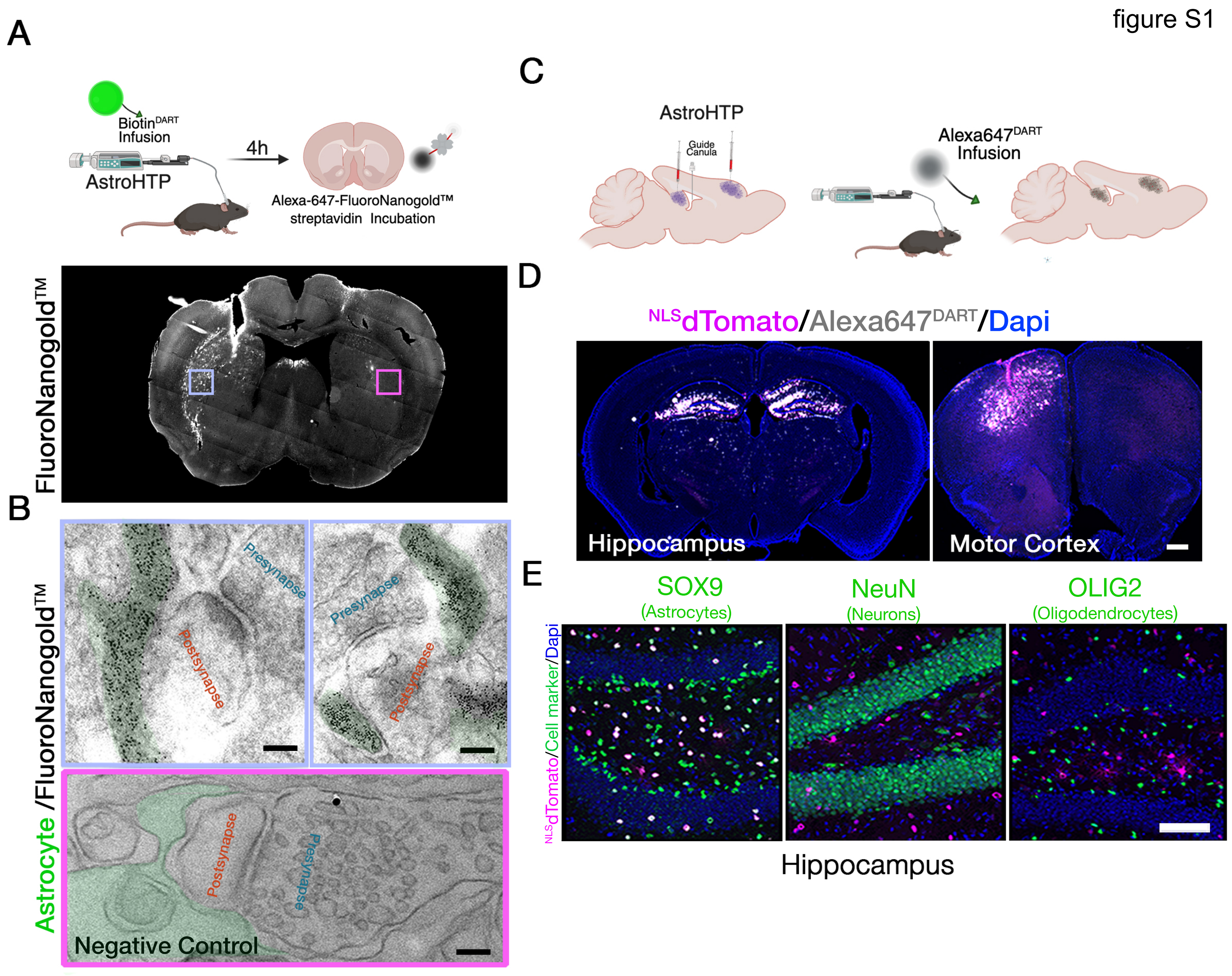

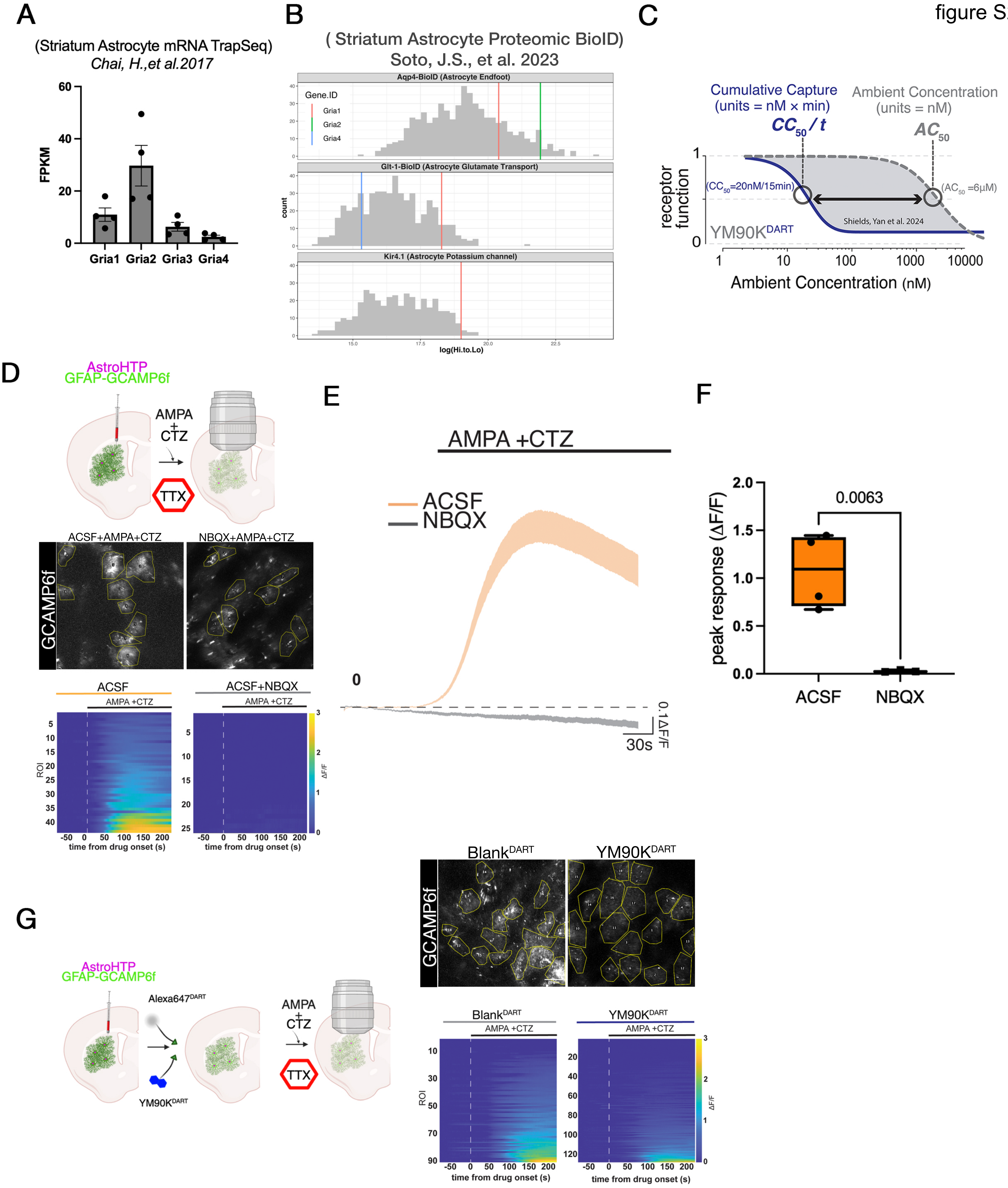

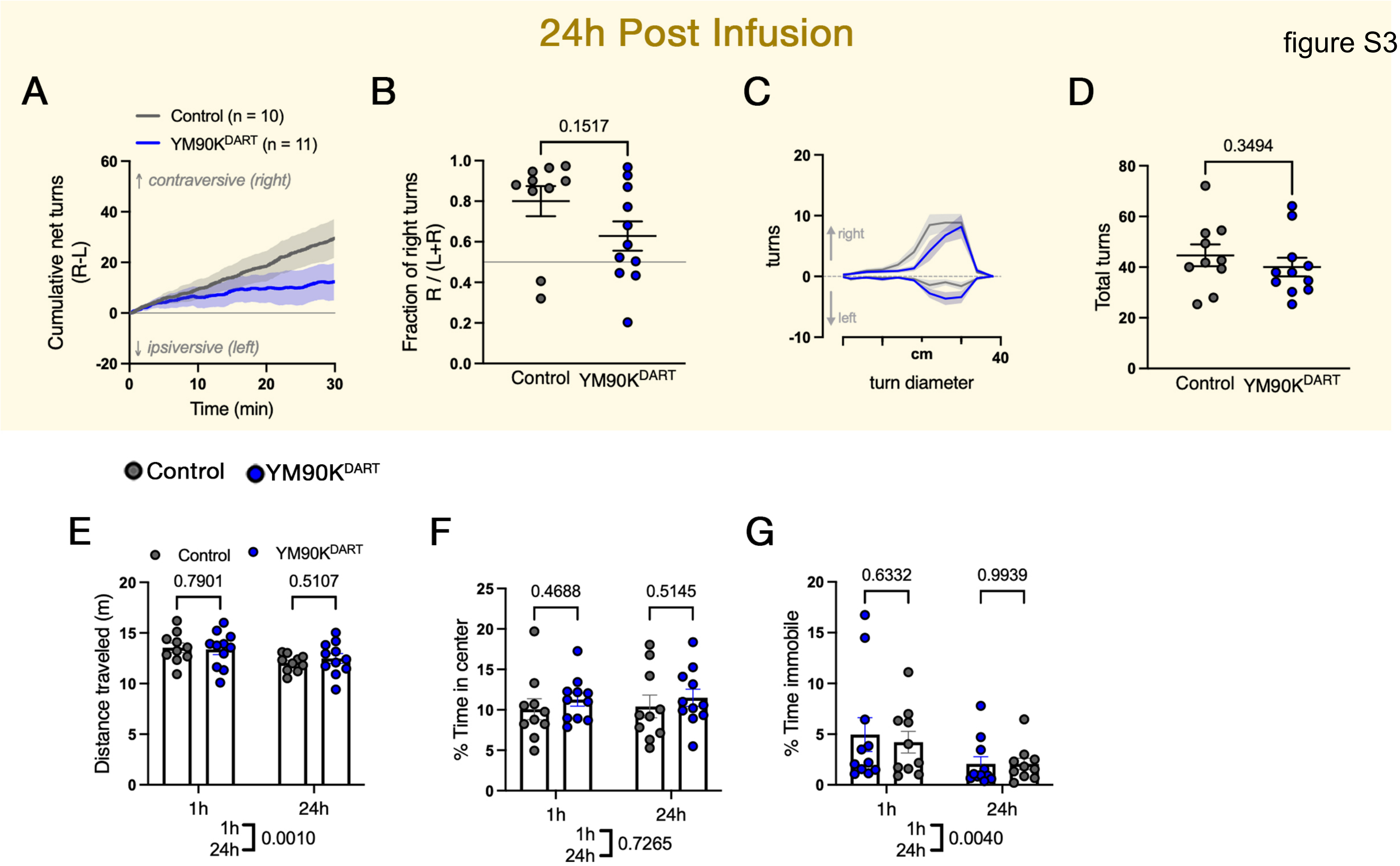

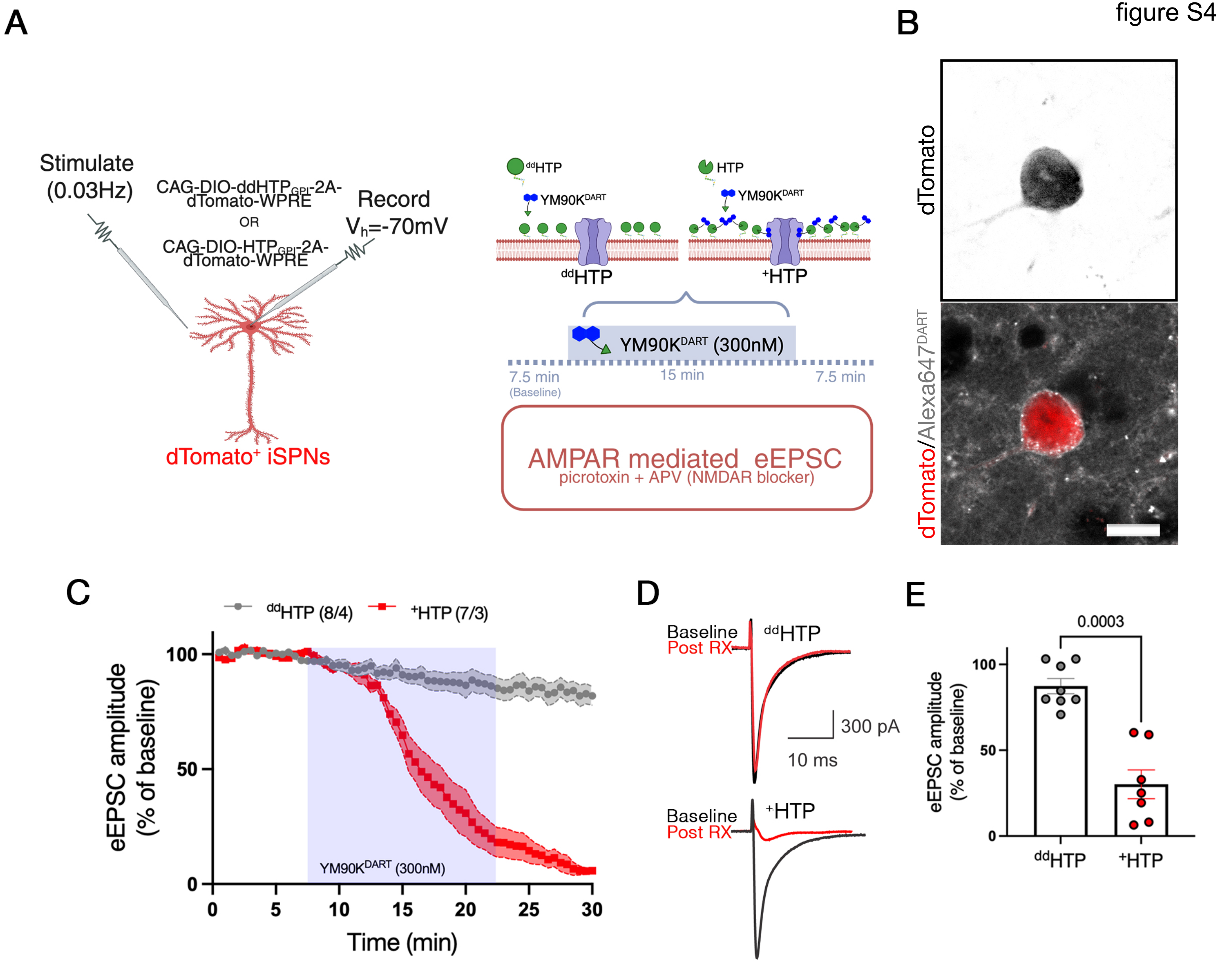

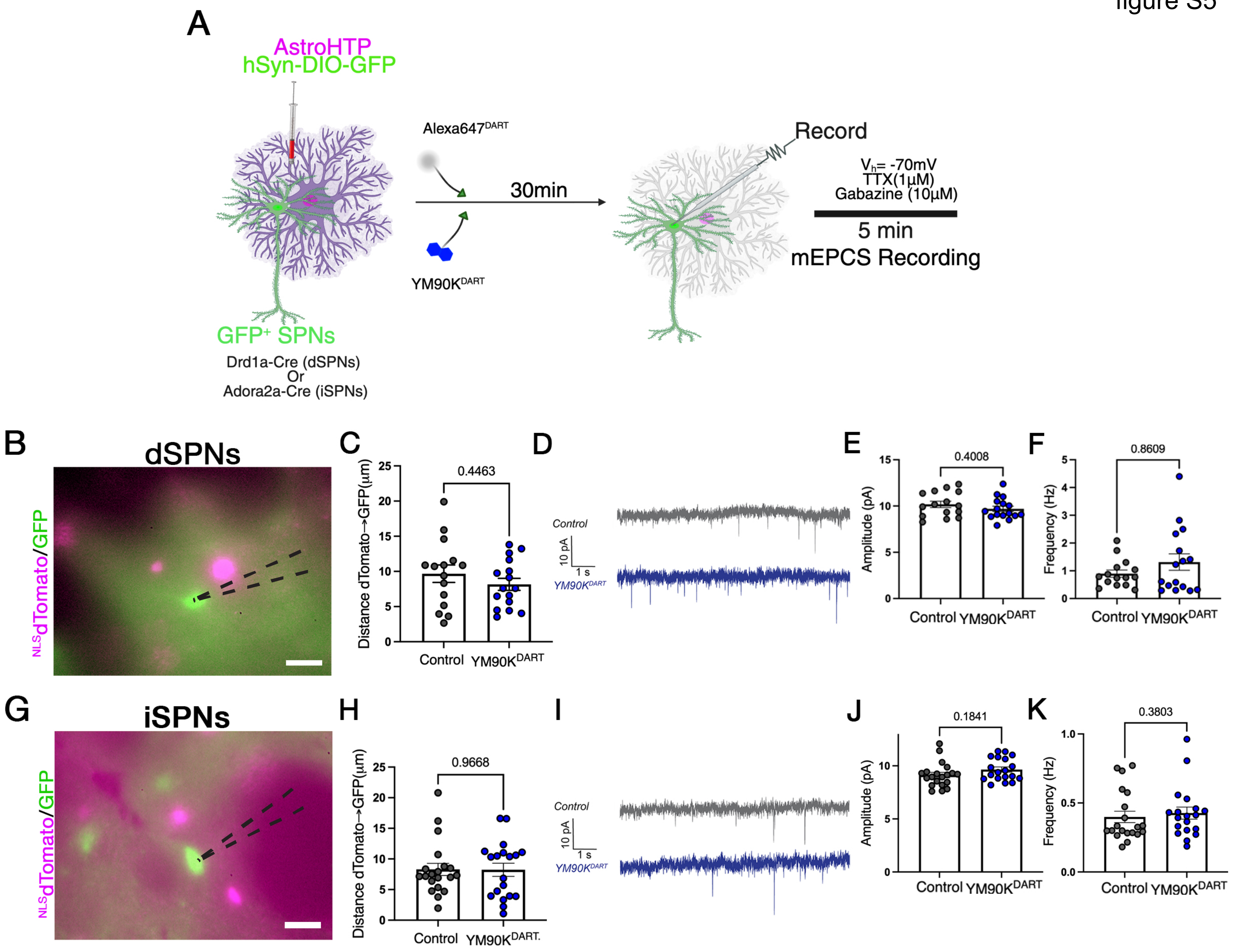

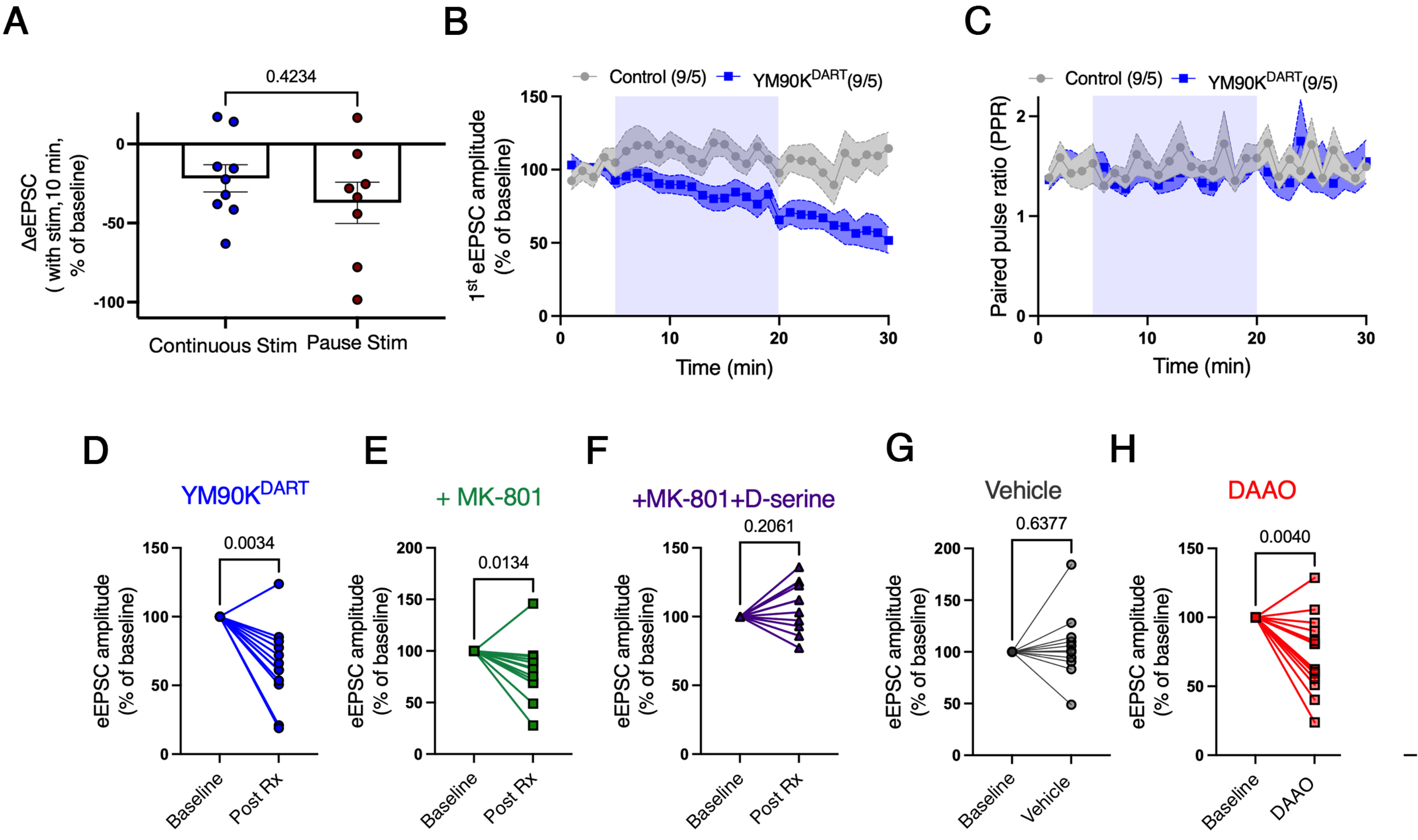
